# Online single-cell data integration through projecting heterogeneous datasets into a common cell-embedding space

**DOI:** 10.1101/2021.04.06.438536

**Authors:** Lei Xiong, Kang Tian, Yuzhe Li, Qiangfeng Cliff Zhang

**Author notes:** Co-first authorship (Lei Xiong, Kang Tian). Correspondence (Q.C.Z.).

## Abstract

Computational tools for integrative analyses of diverse single-cell experiments are facing formidable new challenges including dramatic increases in data scale, sample heterogeneity, and the need to informatively cross-reference new data with foundational datasets. Here, we present SCALEX, a deep-learning method that integrates single-cell data by projecting cells into a batch-invariant, common cell-embedding space in a truly online manner (*i.e.*, without retraining the model). SCALEX substantially outperforms online iNMF and other state-of-the-art non-online integration methods on benchmark single-cell datasets of diverse modalities, (e.g., scRNA-seq, scATAC-seq), especially for datasets with partial overlaps, accurately aligning similar cell populations while retaining true biological differences. We showcase SCALEX’s advantages by constructing continuously expandable single-cell atlases for human, mouse, and COVID-19 patients, each assembled from diverse data sources and growing with every new data. The online data integration capacity and superior performance makes SCALEX particularly appropriate for large-scale single-cell applications to build-upon previously hard-won scientific insights.

## INTRODUCTION

Single-cell experiments enable the decomposition of samples into their constituent, diverse cell-types and cell states^1–4^. Many computational tools have been developed for integrative analysis of single-cell datasets, all seeking to separate biological variations from non-biological noise, such as batch effects of different donors, conditions, and/or analytical platforms^5, 6^. The scope of the integration task is expanding rapidly with technical advances for single-cell studies, which continue to grow larger and larger in scale, now exceeding 1 million cells in some cases^7, 8^. Moreover, the range of examined sample types is also increasing, and datasets now often include highly heterogenous cell subsets^9, 10^. Most importantly, as single-cell studies become more routine, new studies should be informatively cross-referenced to foundational research stuides^7, 8, 11–15^. Thus, there is a growing need for integration tools that can manage single-cell data of large-scale and complex cell-type compositions while also supporting accurate alignment to and exploration within existing datasets.

Most current single-cell data integration methods (e.g., Seurat^16–18^, MNN^19^, Harmony^20^, Conos^21^, Scanorama^22^, BBKNN^23^, *etc.*) are based on the searching across batches for cell-correspondence, for instance similar individual cells or cell anchors/clusters. These methods suffer from three limitations. First, they are prone to mixing cell populations that only exist in some batches, which becomes a severe problem for the integration of complex datasets that contain non-overlapping cell populations in each batch (*i.e.*, partially overlapping data)^16, 17^. Second, they require computational resources that increase dramatically as the number of cells and of batches increase, making these methods increasingly unsuitable for today’s large-scale single-cell datasets^7, 8, 11–15^. Finally, these methods can only remove batch effects from the current dataset being assessed. Each time a new dataset is added, it requires an entirely new integration process that changes the existing integration results of previous studies. This requirement severely limits a tool’s ability to continuously integrate arriving new single-cell data without recalculating existing integrations from scratch, a capacity referred to as “online” data integration^24^.

Online data integration ability is becoming increasingly crucial with today’s single-cell experiments. The recently developed tool, online iNMF^24^, an online version of LIGER^25^, iteratively applies integrative non-negative matrix factorization (iNMF) to decouple the shared and dataset-specific factors related to cell identities, and thus is able to incorporate new data with existing datasets on-the-fly. Another recently developed package, scvi-tools^26^, combining scVI^27^ with scArches^28^, applies a conditional variational autoencoder (VAE)^29^ framework to model the inherent distribution of the input single-cell data for data integration. However, the conditional VAE design of scVI requires model augmentation and retraining when integrating new data, meaning that scVI is not an online method.

Here, we developed SCALEX as a method for online integration of heterogeneous single-cell data based on a VAE framework. The encoder of SCALEX is designed to be a data projection function that only preserves batch-invariant biological data components when projecting single-cells. Importantly, the projection function is a generalized one that requires no retraining on new data, thus allowing SCALEX to integrate single-cell data in an online manner. Working with an extensive collection of benchmark datasets, we demonstrate that SCALEX substantially outperforms online iNMF as well as non-online single-cell data integration tools, in terms of integration accuracy, scalability, and computationally efficiency. The advantages make SCALEX particularly appropriate for the integration and research utilization of today’s single-cell datasets, which continue to grow along with the ongoing explosion of single-cell studies in biology and medicine.

## RESULTS

### SCALEX implements a generalized encoder that enables online integration of single-cell data

To enable online integration, the fundamental design concept underlying SCALEX is to implement a *generalized* projection function that disentangles the batch-related components away from the batch-invariant components of single-cell data and projects the batch-invariant components into a common cell-embedding space. We previously applied VAE and designed SCALE (Single-Cell ATAC-seq Analysis via Latent feature Extraction) to model and analyze single-cell ATAC-seq data^30^. We found that the encoder of SCALE has the potential to disentangle cell-type-related and batch-related features in a low-dimensional embedding space.

Here, to obtain a generalized encoder for data projection without retraining, SCALEX includes three specific design elements (Fig. 1a, Fig. S1, Methods). First, SCALEX implements a batch-free encoder that extracts only biological-related latent features (*z*) from input single-cell data (*x*) and a batch-specific decoder^29^ that reconstructs the original data from *z* by incorporating batch information back during data reconstruction. Supplying batch information only to the decoder focuses the encoder exclusively on learning the batch-invariant biological components, which is crucial for the encoder generalizability. In contrast, scVI includes a set of batch-conditioned parameters into its encoder, which restrains the encoder from the generalizability with new batches and thus precludes online data integration. Second, SCALEX includes a Domain-Specific Batch Normalization (DSBN)^31^ layer using multi-branch Batch Normalization^32^ in its decoder to support incorporation of batch-specific variations during single-cell data reconstruction. Third, the SCALEX encoder employs a mini-batch strategy that samples data from all batches (instead of single batches), which more tightly follows the overall distribution of the input data. Note that each mini-batch is subjected to a Batch Normalization layer in the encoder to adjust the deviation of each mini-batch and to align it to the overall input distribution.

**Fig. 1.**
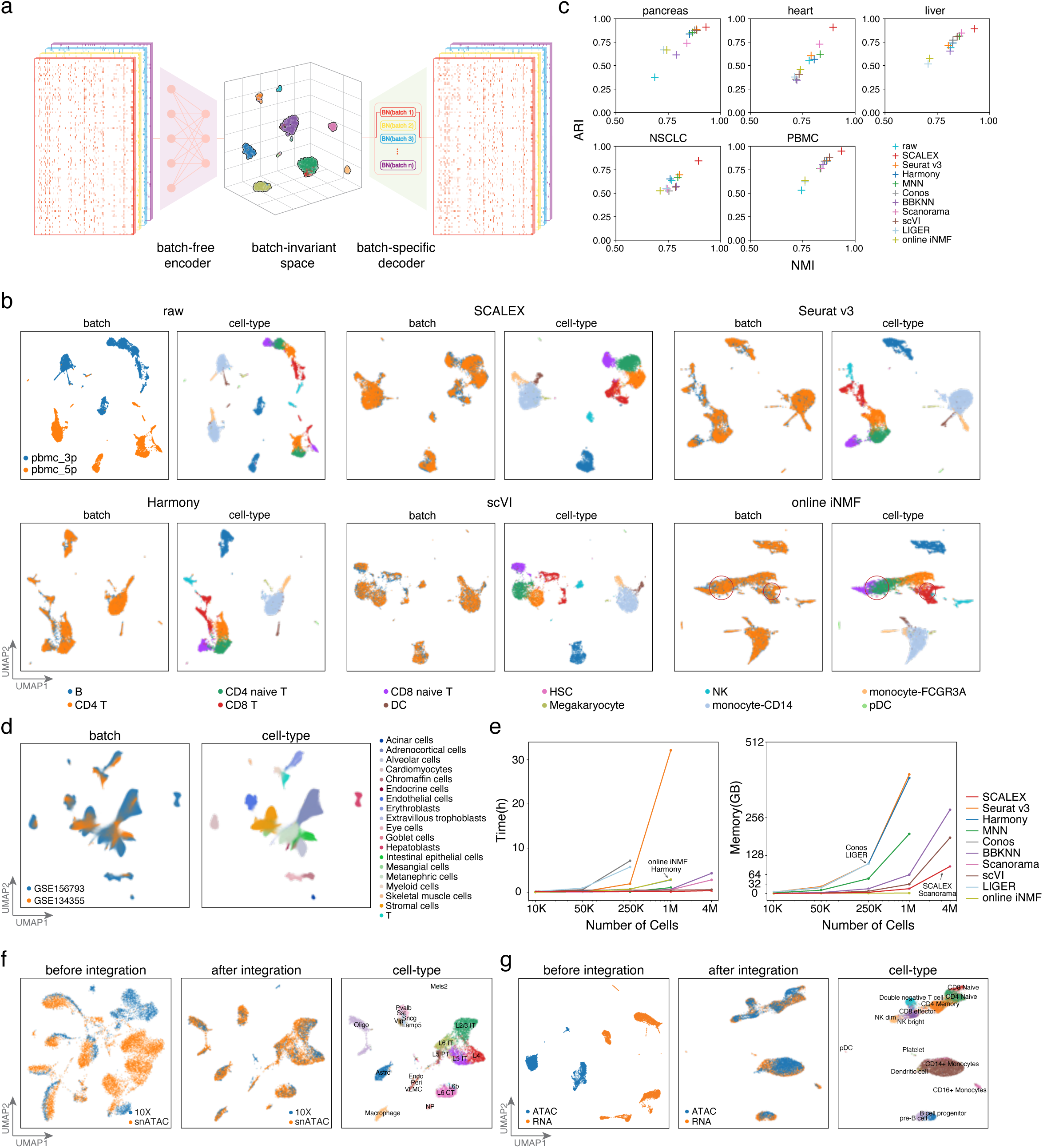
The design and performance of SCALEX for single-cell data integration. **a**, SCALEX models the global structure of single-cell data using a variational autoencoder (VAE) framework. **b**, UMAP embeddings of the *PBMC* dataset before and after integration by indicated methods. Cells are colored by batch (left) and cell-type (right). Misalignments are highlighted with red circles. **c**, Scatter plot comparing SCALEX and the other state-of-the-art single-cell data integration tools in terms of the ARI score (y-axis) and the NMI score (x-axis), based on the Leiden clustering results in the latent space across the indicated benchmark datasets. **d**, UMAP embeddings of the SCALEX integration of the Human Fetal Atlas dataset after integration by SCALEX, colored by batch and cell-type. **e**, Comparison of computation efficiency based on datasets of different sizes sampled from the whole Human Fetal Atlas dataset) including runtime (left) and memory usage (right). Online iNMF was not successfully tested on 4M data due to a HDF5 file conversion issue for large data (Methods). **f**, UMAP embeddings of the mouse brain scATAC-seq dataset before (left) and after integration (middle, right); colored by data batch or cell-types. **g**, UMAP embeddings of the PBMC scRNA-seq and scATAC-seq cross-modality dataset before (left) and after SCALEX integration (middle, right); colored by batch or cell-type.

We conducted extensive analyses of SCALEX hyperparameters and also tested the specific contributions of each design element by implementing a set of SCALEX test-variants, each lacking an individual or a combination of the design elements, and evaluating their performance for single-cell data integration (Supplementary Note). We found that each design element is crucial for the integration performance of SCALEX. More importantly, the combination of these design elements renders the encoder of SCALEX a generalized function capable of accurate projection of single cell data from different batches into a batch-invariant cell-embedding space, making SCALEX a truly online data integration method.

### SCALEX integration is substantially more accurate than state-of-the-art single-cell data integration methods

We extensively assessed the basic data integration performance of SCALEX (Methods), following the evaluative framework proposed in a recent comparative study^33^. We examined multiple well-curated scRNA-seq datasets, including human *pancreas* (eight batches of five studies)^34–38^, *heart* (two batches of one study)^39^ and *liver* (two studies)^40, 41^; as well as human non-small-cell lung cancer (*NSCLC*, four studies)^42–45^ and peripheral blood mononuclear cells (*PBMC*; two batches assayed by two different protocols)^16^. Our comparison included online iNMF and other state-of-the-art non-online single-cell data integration methods, including Seurat v3, Harmony, MNN, Conos, BBKNN, Scanorama, LIGER (*i.e.*, batch iNMF), and scVI. We evaluated the integration performance of these tools based on the benchmark datasets by Uniform Manifold Approximation and Projection (UMAP)^46^ embedding visualization as well as a series of scoring metrics ^19, 20, 47–49^.

With UMAP embedding, we note that all of the raw datasets displayed strong batch effects, with cell-types that were common in different batches separately distributed. Overall, SCALEX, Seurat v3, and Harmony achieved the best integration performance for most of the datasets by merging common cell-types across batches while keeping disparate cell-types apart (Fig. S2). MNN, scVI, and Conos integrated many datasets but left some common cell-types not well-aligned. Online iNMF, LIGER, BBKNN, and Scanorama often had unmerged common cell-types, and sometimes incorrectly mixed distinct cell-types together. For example, considering the T cell populations between the two batches in the *PMBC* dataset (Fig. 1b), while SCALEX, Seurat v3, Harmony, MNN, scVI integrations were effective, online iNMF misaligned some of the CD4 naïve T cells with CD8 naïve T cells, and misaligned some NK cells with CD8 T cells.

SCALEX substantially outperformed all of the other methods for cell-type clustering, as assessed by the adjusted Rand Index (ARI)^47^ and the Normalized Mutual Information (NMI)^48^ (Fig. 1c, Fig. S3-S4). To quantify cell-type separation and batch mixing we used two paired metrics (Methods): a pair comprising the Silhouette score^49^ and the batch entropy mixing score^19^, as well as a pair comprising the cell-type and integration local inverse Simpson’s Indexes (cLISI and iLISI)^20^. Overall, SCALEX achieved the highest scores for cell-type separation, and tied with Seurat v3 and Harmony as the best-performing methods on the batch mixing metrics (Fig. S4). Interestingly, we observed that both LIGER and online iNMF often scored the lowest for cell-type separation yet the highest for batch mixing. However, after careful investigation, we concluded that a higher batch mixing score does not necessarily indicate better data integration, but instead often indicates an issue of over-correction, which we consider in-depth in a dedicated subsection below.

### SCALEX is scalable to Atlas-level datasets and accommodates diverse data modalities

Single-cell datasets that contain a large number of cells and consist of heterogenous and complex samples from multiple tissues have been termed “Atlas-level” datasets in a recent comparative study^33^. These Atlas-level datasets are posing new challenges to data integration tools. We tested the scalability and computation efficiency of SCALEX by applying it to a typical Atlas-level dataset, the Human Fetal Atlas dataset, which contains 4,317,246 cells from two data batches, GSE156793 and GSE134355 (Methods)^8, 15^ (Fig. S5a,b). SCALEX accurately integrated these two batches, showing good alignment of the same cell-types (Fig. 1d). In addition to SCALEX, only BBKNN, Scanorama, and scVI can be used to integrate this Atlas-level dataset, however, their integrations does not separate and align the cell-types well, as indicated by the UMAP embeddings (Fig. S5c) and the low cell-type separation and batch mixing scores (Fig. S5d).

We compared the computational efficiency of different methods using down-sampled datasets (of 10 K, 50 K, 250 K, 1 M, and 4M cells) from this Human Fetal Atlas dataset. Both SCALEX and online iNMF consumed very efficient runtime and memory that increased only linearly with data size. scVI also is scalable to 4M cells with acceptable memory usage, whereas Seurat v3, Harmony, Conos, and LIGER consumed runtime and/or memory that increased exponentially, thus did not scale beyond 1M cells on a workstation of 64 CPU cores and 256 GB memory (Fig. 1e). Notably, the deep learning framework of SCALEX enables it to run very efficiently on GPU devices, requiring much reduced runtime (using about 20 minutes and 90 GB of memory on the 4M dataset).

SCALEX can be used to integrate other modalities of single-cell data (*e.g.*, scATAC-seq^50, 51^, CITE-Seq^52^, *etc.*) and cross-modality data (*e.g.*, simultaneous analysis of scRNA-seq and scATAC-seq) (Methods). SCALEX substantially outperformed all other methods for integration of mouse brain scATAC-seq datasets (two batches assayed by snATAC and 10X)^53^ (Fig. 1f, Fig. S6a-c), and performed well for integration of additional single-cell data modalities including CITE-seq^52^ and spatial transcriptome MERFISH data^54^ (Fig. S6d,e). We also used SCALEX to integrate a cross-modality dataset (scRNA-seq and scATAC-seq)^55, 56^ and found that SCALEX correctly integrated the two modalities of data and distinguished rare cells that are specific to the scRNA-seq data, including pDC and platelet cells (Fig. 1g), doing so better than other methods according to both UMAP embeddings and multiple analytical metrics (Fig. S7).

### SCALEX integrates partially overlapping datasets without over-correction

Many recent single-cell datasets, especially Atlas-level datasets, feature high sample heterogeneity and complex cell-type compositions^9, 10^. These datasets often contain partially overlapping batches where each batch contains some non-overlapping cell populations. For example, the *liver* dataset is a partially overlapping dataset where the hepatocyte population contains multiple subtypes specific to different batches: three subtypes are specific to LIVER_GSE124395, and two other subtypes only appear in LIVER_GSE115469 (Fig. S8).

This partial overlap problem presents a major challenge for single-cell data integration and often leads to an issue of over-correction (*i.e.*, mixing of distinct cell-types), especially for those local cell similarity-based methods^16, 17^. For example, Seurat v3 mixed the hepatocyte-CXCL1, hepatocyte-CYP2A13, and hepatocyte-TAT-AS1 cells and Harmony mixed the hepatocyte-CYP2A13 and hepatocyte-TAT-AS1 cells (Fig. 2a). As a global integration method that projects cells into a common cell-embedding space, SCALEX is expected to be less sensitive to this problem. Indeed, we noticed that SCALEX correctly maintained the five hepatocyte subtypes apart (as did scVI. Fig. 2a). Unexpectedly, despite being a global method, online iNMF severely suffered from over-correction, mixing all five hepatocyte subtypes, and even mixing B cells and NK cells (Fig. 2a), presumably because its matrix factoring algorithm forced the alignment of distinct cell-types.

**Fig. 2.**
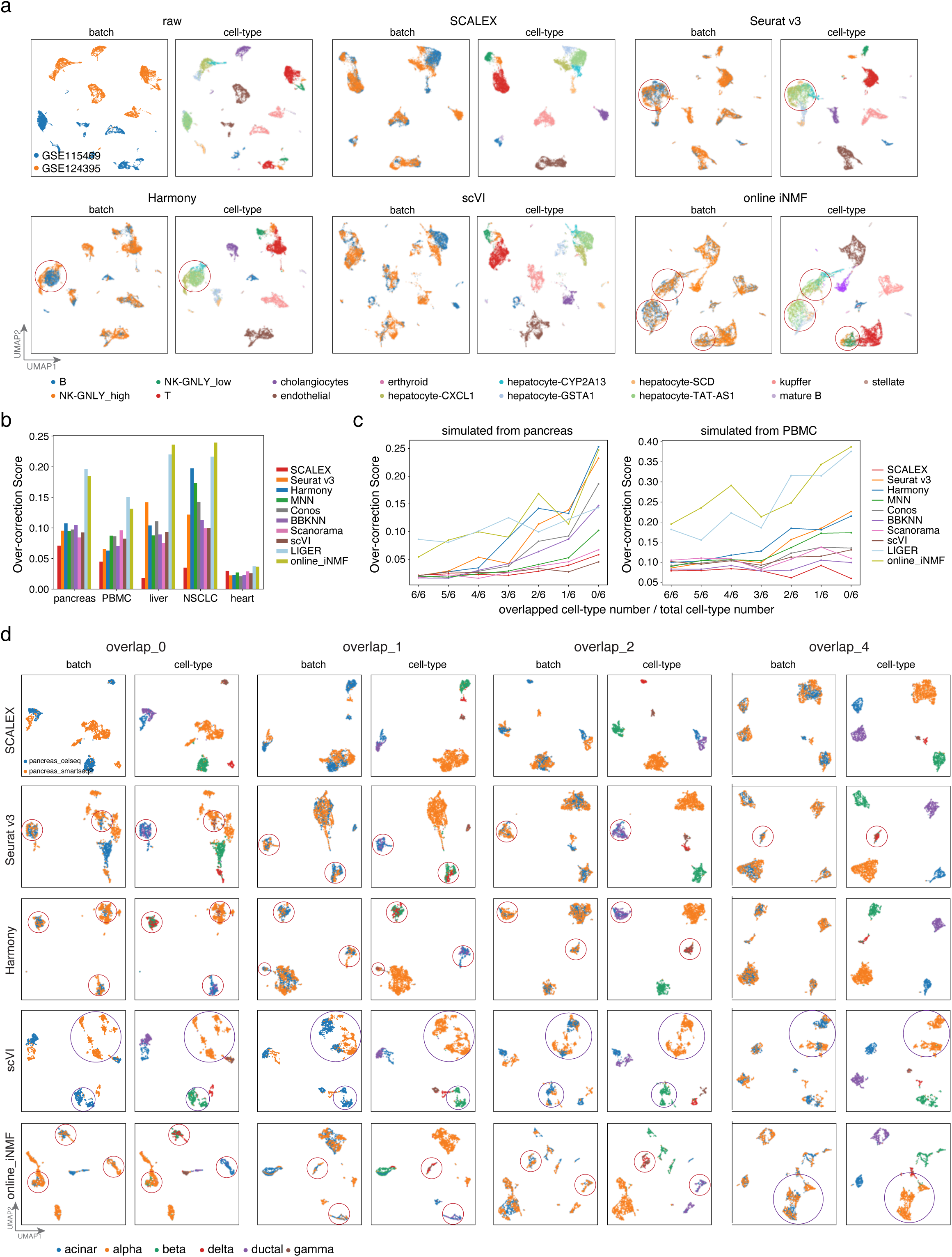
Comparison of integration performance over partially overlapping datasets by different methods. **a**, UMAP embeddings of the integration results by the indicated methods based on the *liver* dataset. **b**, Over-correction score of different methods based on the indicated benchmark datasets. **c**, Over-correction score of the indicated methods based on the simulated datasets, with decreased numbers of common cell-types (obtained by down-sampling the *pancreas* and *PBMC* dataset). **d**, UMAP embeddings of the integration results by the indicated methods based on the simulated *panceas* datasets with different numbers of specified common cell-types. Over-corrections are highlighted with red circles and error-splittings with purple circles, respectively.

We defined an over-correction score, a metric to measure this over-correction problem based on the percent of cells with inconsistent cell-types in the neighborhood for each cell (Methods). Formally, the over-correction score is a negative index, *i.e.*, the higher the over-correction score, the more severe the extent of inaccurate mixing of cell-types. For the benchmark datasets, SCALEX had the lowest over-correction scores (Fig. 2b), whereas online iNMF yielded extremely high over-correction scores.

To systematically characterize the performance of different methods on partially overlapping datasets, we constructed test datasets with a range of common cell-types, that we generated based on down-sampling of the six major cell-types in the *pancreas* dataset (Methods). SCALEX integration was accurate for all cases, aligning the same cell-types without over-correction, whereas Seurat v3, Harmony, and online iNMF frequently mixed distinct cell-types (Fig. 2c-d). Although scVI showed one of the lowest levels of over-correction when integrating partially overlapping datasets, it is prone to mistakenly splitting one cell-type into many small groups. We noted that the severity of over-correction and error-splitting is amplified as the overlapping number decreases (Fig. S9). When there were no common cell-types, both Seurat v3 and Harmony collapsed the six cell-types into three, mixing alpha with gamma cells, beta with delta cells, and acinar with ductal cells to varying extents, whereas scVI split alpha cells into 6 groups. We repeated this down-sampling analysis from the 12 cell-types in the *PBMC* dataset and observed similar results of over-correction and error-splitting (Fig. S10).

### SCALEX increases the scope and resolution of an existing cell space by adding new data through online projection

The generalizability of SCALEX’s encoder to project cells from various sources into a common cell-embedding space without model retraining allows SCALEX to integrate new single-cell data with existing data in an online manner. We tested the online data integration performance of SCALEX for newly arriving data based on the *pancreas* dataset. Prior to projection, we first used SCALEX to integrate the *pancreas* dataset and this accurately removed the strong batch effect that was evident in the raw data (Fig. 3a, Fig. S11a-b).

**Fig. 3.**
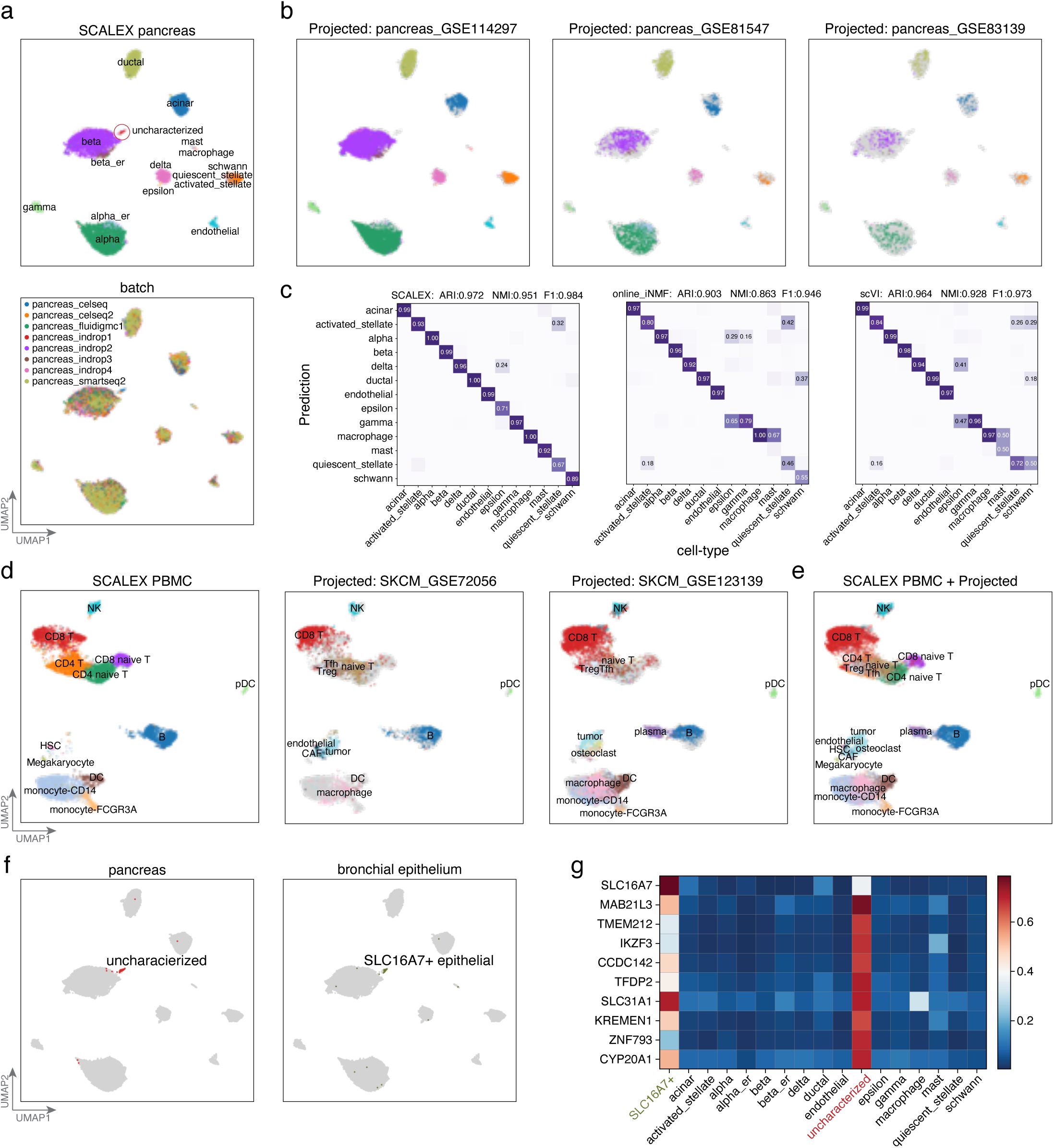
Projecting heterogenous data into a common cell-embedding space. **a**, UMAP embeddings of the *pancreas* dataset after integration by SCALEX, colored by cell-type and by batch. **b**, UMAP embeddings of the common cell space obtained by using SCALEX to project three additional indicated *pancreas* data batches onto the *pancreas* dataset. Cells are colored by cell-type with light gray shadows representing the original *pancreas* dataset. **c**, Confusion matrix between ground truth cell-types and those annotated by different methods. ARI, NMI and F1 scores (top) measure the annotation accuracy. **d**, UMAP embeddings of the common space obtained by using SCALEX to project the two projected melanoma data batches onto the *PBMC* dataset, colored by cell-types with light gray shadows represent the original *PBMC* dataset. **e**, UMAP embeddings of the common cell space that includes the original *PBMC* dataset and the two projected melanoma data batches. **f**, Annotating an uncharacterized small cell population in the *pancreas* dataset by projection of the bronchial epithelium data batches into the pancreas cell space. Only the uncharacterized cells in the *pancreas* dataset (left) and the SLC16A7+ epithelial cells in the bronchial epithelium data batches (right) are colored. **g**, Heatmap showing the normalized expression of the top-10 ranking specific genes for the uncharacterized cell population in different cell-types.

We subsequently projected three new batches of scRNA-seq data^57–59^ for pancreas tissues (Fig. 3b) into this “pancreas cell space” using the same SCALEX encoder trained on the original *pancreas* dataset. After projection, most of the cells in the new batches were accurately aligned to the correct cell-types in the pancreas cell space, enabling their accurate annotation by cell-type label transfer (Fig. 3c, Method). We benchmarked projection accuracy by calculating the ARI, the NMI, and the F1 score to evaluate cell-type annotation by label transfer with cell-type information in the original studies (Methods). We compared the results with online iNMF and scVI, the only two tools that are able to project cells into an *existing* cell-space (note that data projection of scVI needs model retaining through scArches). SCALEX achieved the highest projection accuracy in comparisons with online iNMF and scVI (Fig. 3c). scVI also achieved high accuracy, projecting most cells onto right locations, with only a few exceptions of alpha and ductal cells (Fig. S11c). Online iNMF mixed distinct cell-types when incorporating new batches, *e.g*. projecting some alpha cells onto the locations of gamma and delta cells, and delta cells onto beta cells (Fig. S11c), which in turn led to wrong annotations during label transfer (Fig. 3c).

The ability to project new single-cell data into an existing cell-embedding space allows SCALEX to readily enrich (*i.e.*, to add biological resolution) this cell space with additional informative details. To verify this, we projected two additional melanoma data batches (SKCM_GSE72056, SKCM_GSE123139)^10, 60^ onto the previously constructed PBMC space. Again, SCALEX correctly projected all common cell-types were onto the same locations in the PBMC cell space (Fig. 3d), but online iNMF mixed tumor cells with plasma, monocyte and CD8 T cells, and scVI split the CD8T cells into several distinct groups (Fig. S12). Importantly, we noticed that for the tumor and plasma cells only present in the melanoma data batches, SCALEX did not project these cells onto any existing cell populations in the PBMC space; rather, it projected them onto new locations close to similar cells, with the plasma cells projected to a location near B cells, and the tumor cells projected to a location near HSC cells (Fig. 3e). This indicates that SCALEX can enrich an existing cell space with new cell-types through data projection.

SCALEX projection also enables *post hoc* annotation of unknown cell-types in an existing cell space using new data. For instance, we noted a group of previously uncharacterized cells in the *pancreas* dataset (Fig. 3a). We found that these cells displayed high expression levels of known epithelial gene markers (Methods). We therefore assembled a collection of epithelial cells from the *bronchial epithelium* dataset^61^, and then projected these epithelial cells onto the pancreas cell space. We found that a group of antigen-presenting airway epithelial (SLC16A7+ epithelial) cells were projected onto the same location of the uncharacterized cells (Fig. 3f). These data, together with the observation that both cell populations showed similar marker gene expression (Fig. 3g), suggest that these uncharacterized cells are also SLC16A7+ epithelial cells. Note that online iNMF and scVI were not able to identify this small group of epithelial cells, because they were split into several smaller groups and/or were often mixed with other cell-types (Fig. S2). SCALEX thus enables discovery science in cell biology by supporting exploratory analysis with large numbers of diverse datasets.

### SCALEX integration constructs expandable single-cell atlases

The ability to combine heterogenous data into a common cell-embedding space makes SCALEX a powerful tool to construct a single-cell atlas from a collection of diverse datasets. We applied SCALEX integration to three large and complex datasets: the Mouse Atlas dataset (comprising multiple organs from two studies assayed by 10X, Smart-seq2, and Microwell-seq^12, 14^), the Human Atlas dataset (comprising multiple organs from two studies assayed by 10X and Microwell-seq^15, 62^), and the Human Fetal Atlas dataset^8, 15^ (Fig. S13).

Despite the strong batch effects in the raw data, SCALEX accurately integrated the three batches of the Mouse Atlas data into a common cell-embedding space (Fig. 4a-c, Fig. S14a). Common cell-types were well-aligned at the same position in the cell space, including B, T, and endothelial cells presented in all tissues, and proximal tubule, urothelial, and hepatocytic cells from particular tissues. Distinct cell-types were located separately, such as sperm, Leydig, and small intestine cells from the Microwell-seq data, keratinocyte stem cells and large intestine cells from the Smart-seq2 data, indicating that biological variations were well preserved (Fig S14b,c). We compared SCALEX with all other methods and found that SCALEX performed the best for cell-type clustering, especially for avoiding over-correction (Fig. 4d,e, Fig. S13b).

**Fig. 4.**
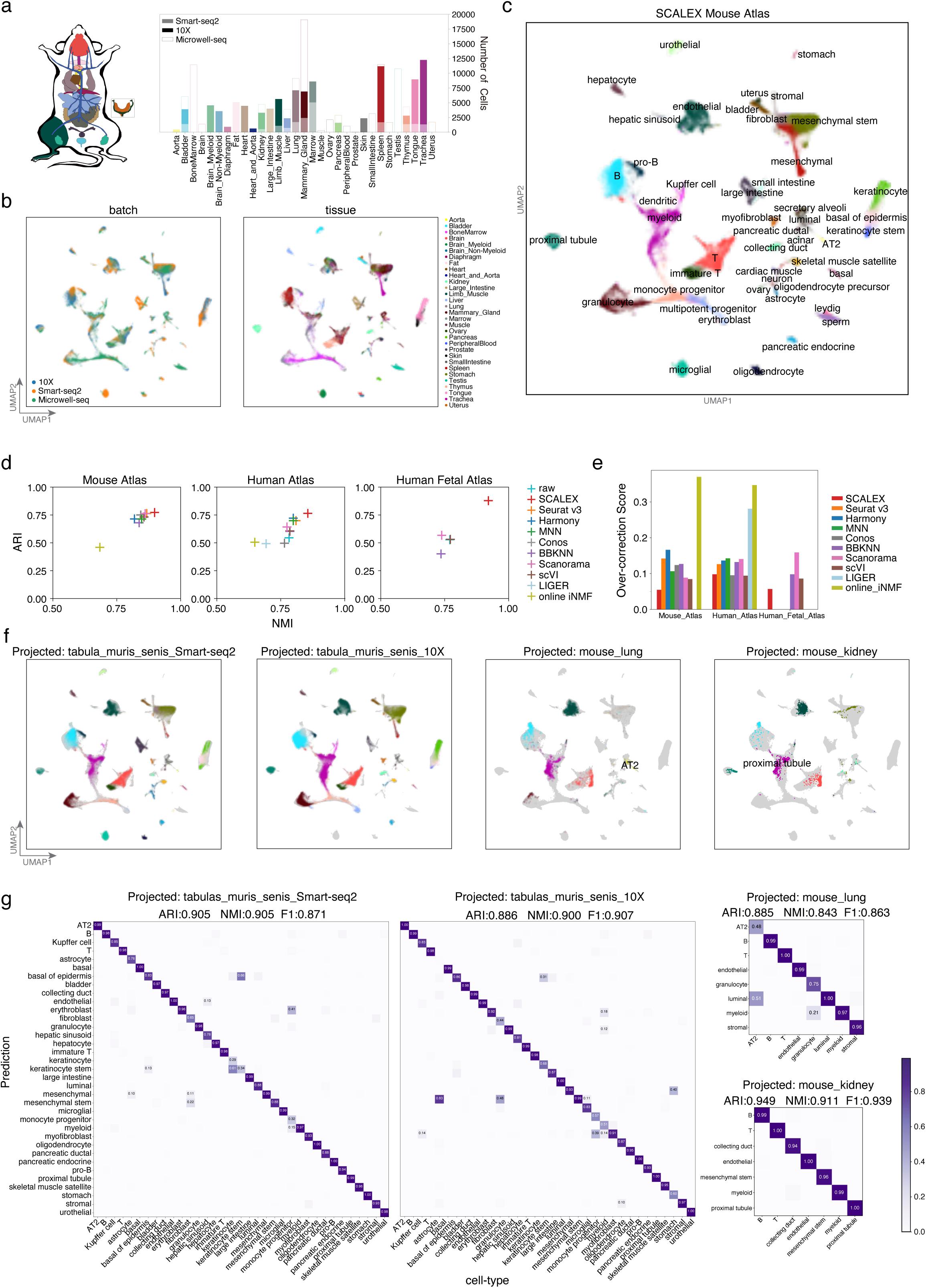
Construction of an expandable mouse single-cell atlas. **a,** Datasets acquired using different technologies (Smart-seq2, 10X, and Microwell-seq) covering various tissues used for construction of the mouse atlas. **b**, UMAP embeddings of the Mouse Atlas dataset colored by batch and tissue. **c**, UMAP embeddings of the Mouse Atlas after SCALEX integration, colored by cell-type. **d**, Scatter plot showing a quantitative comparison of the ARI score (y-axis) and the NMI score (x-axis) based-on the Leiden clustering results on the latent space based on the Human Atlas, Mouse Atlas, and Human Fetal Atlas datasets. **e**, Comparison of over-correction score by the indicated methods based on the Human Atlas, Mouse Atlas, and Human Fetal Atlas datasets. **f**, UMAP embeddings of the common cell space obtained by using SCALEX to project the two *Tabula Muris Senis* data batches and two mouse tissues (lung and kidney) data batches onto the Mouse Atlas dataset. Cells are colored by cell-type with light gray shadows representing the original Mouse Atlas dataset. **g**, Confusion matrix of the cell-type annotations by SCALEX and those in the original studies. Color bar represents the percentage of cells in confusion matrix *C_ij_* known to be cell-type *i* and predicted to be cell-type *j*.

Importantly, atlases generated with SCALEX can be further expanded by projecting new single-cell data to support comparative studies of cells both in the original atlas and in the new data. To illustrate this utility, we projected two additional data batches of aged mouse tissues from *Tabula Muris Senis* (Smart-seq2 and 10X)^13^ and two single tissue datasets (lung and kidney)^63^ onto the SCALEX Mouse Atlas cell space. We found that cells in the new data batches were correctly projected onto the locations of the same cell-types in the cell-embedding space of the initial atlas (Fig. 4f) as confirmed by the accurate cell-type annotations for the new data by label transfer (Fig. 4g. Methods).

Following the same strategy, we constructed a SCALEX Human Atlas by integration of multiple tissues from two studies (GSE134355, GSE159929) (Fig. S15a,b). SCALEX effectively eliminated the batch effects in the original data and integrated the two datasets (Fig. S15c,d). Again, we were able to correctly project two additional human skin datasets (GSE130973, GSE147424)^64, 65^ onto the Human Atlas cell space (Fig. S15e), and accurately annotated these projected skin cells (Fig. S15f. Methods). In sum, these results illustrate that SCALEX enables: i) researchers to evaluate their project-specific single cell datasets by leveraging existing information in large-scale (and ostensibly well annotated) cell atlases; and ii) atlas creators to informatively integrate new datasets and derive new biological insights from new research programs.

### An integrative SCALEX COVID-19 PBMC Atlas revealed distinct immune responses among COVID-19 patients

Many single-cell studies have been conducted to analyze COVID-19 patient immune responses^66–73^. However, these studies often suffer from small sample size and/or limited sampling of various disease states^67, 73^. For a comprehensive study, we used SCALEX to generate a COVID-19 PBMC Atlas, integrating data from nine COVID-19 studies, involving a total of 860,746 single cells in 10 batches^66–72^ (Fig. 5a, Fig. S16a,b). We identified 22 cell-types, each of which has support from gene expression data for canonical markers (Fig. 5b,c, Fig. S16c,d. Methods). Cells across different studies were integrated accurately with the same cell-types aligned together, confirming the integration performance of SCALEX (Fig. 5c, Fig. S16d), which was much better than the other methods in terms of cell-type clustering (Fig. S16a,b).

**Fig. 5.**
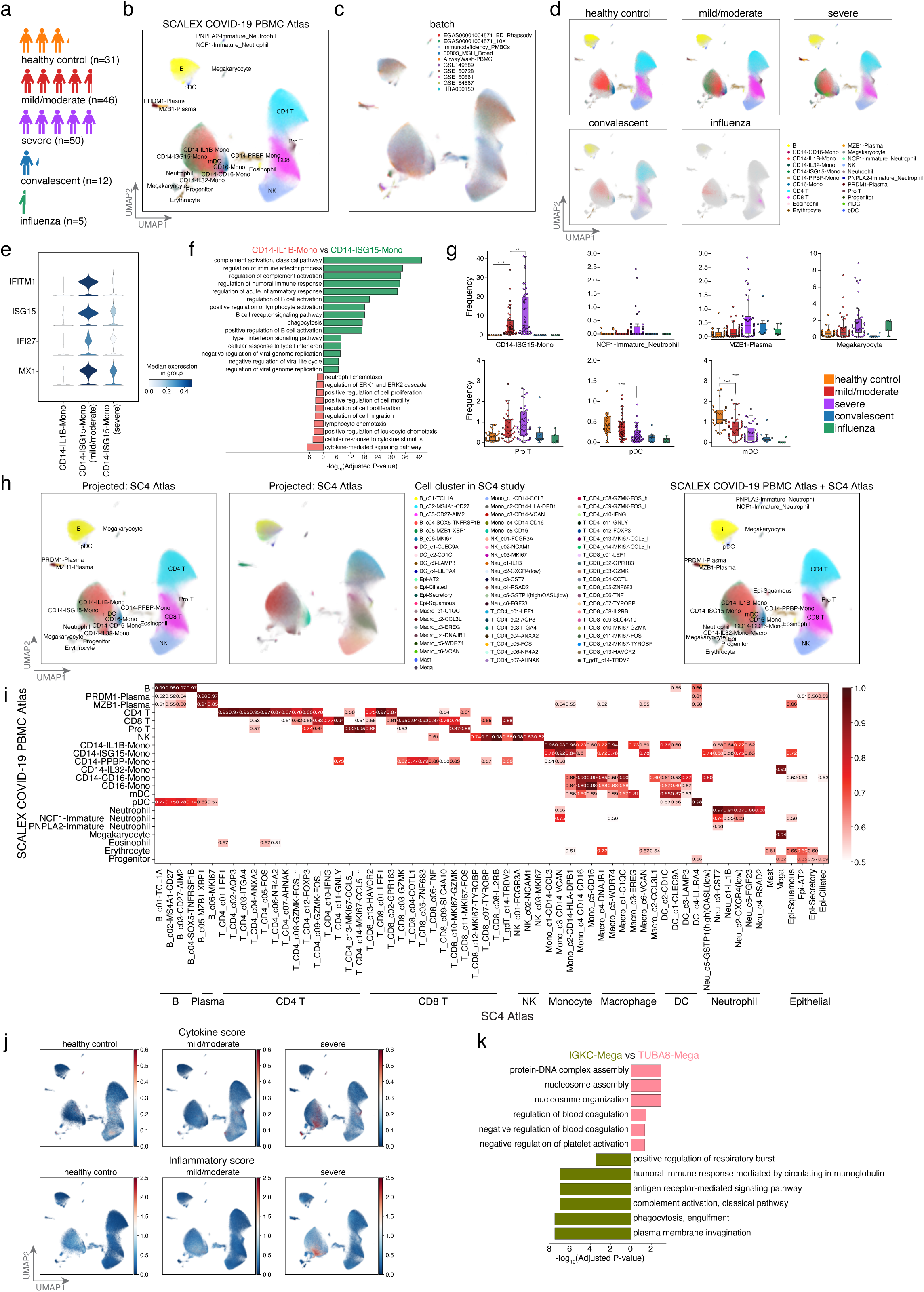
Online integration of COVID-19 PBMC Atlas. **a**, The COVID-19 PBMC Atlas dataset composition, including healthy controls and influenza patients, as well as mild/moderate, severe, and convalescent COVID-19 patients. **b**,**c** UMAP embeddings of the COVID-19 PBMC Atlas after SCLAEX integration colored by cell-type (**b**), and by batch (**c**). **d**, UMAP embeddings of the COVID-19 PBMC Atlas separated by disease state. Cells are colored by cell-type with light gray shadows representing the other disease state cells. **e**, Stacked violin plot of differentially-expressed ISGs among CD14 monocytes across disease states. **f**, GO terms enriched in the differentially-expressed genes for CD14-IL1B-Mono and CD14-ISG15-Mono cells. **g**, Cell-type frequency across healthy and influenza controls, and among mild/moderate, severe, and convalescent COVID-19 patients. Dirichlet-multinomial regression was used for pairwise comparisons, ***p<0.001, **p<0.01, *p<0.05. **h**, UMAP embeddings of the common cell space obtained by using SCALEX to project the SC4 Atlas (Single Cell Consortium for COVID-19 in China). Cells are colored by cell-type from label transfer based-on the locations in the COVID-19 PBMC Atla*s* dataset (left) and cell clusters in original SC4 study (middle), and Unified UMAP embeddings combining the SCALEX COVID-19 PBMC Atlas and the SC4 Atlas (right). **i**, Similarity matrix of meta-cell representations for cell-types between the SCALEX COVID-19 PBMC Atlas and SC4 in the common cell-embedding space after SCALEX projection. Color bar represents the Pearson correlation coefficient between the average meta-cell representation of two cell-types from a respective data batch. **j**, UMAP embeddings of the SCALEX COVID-19 PBMC Atlas colored by the cytokine score and the inflammatory score. **k**, GO terms enriched in the differentially-expressed genes for TUBA8-Mega and IGKC-Mega cells.

Interestingly, we found that some cell subpopulations were differentially associated with patient status (Fig 5d). A subpopulation of CD14 monocytes (CD14-ISG15-Mono) was characterized by its high expression of Type I interferon-stimulated genes (ISGs) and genes enriched with immune-response-related GO terms (Fig 5e,f). The frequency of CD14-ISG15-Mono cells increased significantly from mild/moderate to severe patients (Fig. 5g, Fig. S17a. Methods). Within the COVID-19 patients, we observed a significant decrease in ISG gene expression in CD14-ISG15-Mono cells between the mild/moderate and severe cases, suggesting an immune exhaustion-like response in severe COVID-19 patients (Fig. 5e).

Additionally, a neutrophil subpopulation (NCF1-Immature_Neutrophil), characterized by decreased expression of the genes responsible for neutrophil activation but elevated expression of genes enriched with viral-process-related GO terms, was specifically enriched in severe verse mild/moderate patients (Fig. S17b,c). A plasma cell subpopulation (MZB1-Plasma), characterized by decreased expression of genes related to antibody production and enriched for immune and inflammatory response-related GO terms, were also enriched in severe patients (Fig. S17d,e). Thus, the SCALEX COVID-19 PBMC atlas, generated by integrating a highly diverse collection of single-cell data from individual studies, identified multiple immune cell-types that become progressively dysfunctional during COVID-19 disease progression. Importantly, these cell trends were not and could not have been detected in the small-scale, individual studies that served as the basis for our SCALEX COVID-19 PBMC atlas.

### Online integration of the SCALEX COVID-19 PBMC Atlas with the SC4 consortium study

Our analysis based on the SCALEX COVID-19 PBMC Atlas yielded findings consistent with two conclusions from the Single Cell Consortium for COVID-19 in China (SC4) study, a recent large-scale effort that generated a single-cell atlas of over 1 million cells from 171 COVID-19 patients and 25 healthy controls^7^ (Fig. S18a). First, both studies observed the same set of immune cell subpopulations which displayed differential associations with COVID-19 severity. The proportions of CD14 monocytes, megakaryocytes, plasma cells, and pro T cells were elevated with increasing disease severity, while the proportion of pDC and mDC cells decreased (Fig. 5g). Second, based on calculating the same cytokine score and inflammatory score (defined in the SC4 study) for the cells in our SCALEX COVID-19 PBMC Atlas, we confirmed that the megakaryocytes and monocyte subpopulations are associated with cytokine storms triggered by SARS-Cov2 infection and are further elevated in severe patients^74^ (Fig. 5j. Methods).

SCALEX’s online integration capacity enables us to project the SC4 consortium dataset into the cell space of the SCALEX COVID-19 PBMC Atlas. We found that the cell-types of two atlases were well-aligned (Fig. 5h,i, Fig. S18b,c). Integration of the SC4 data further substantially improved both the scope and resolution of the SCALEX COVID-19 PBMC Atlas. First, this data added macrophages and epithelial cells to the cell space, enabling investigation of their potential involvement in COVID-19. The integration also supported more precise characterization of specific cell subpopulations. For example, the megakaryocyte population, not distinguished in either the SCALEX COVID-19 PBMC Atlas or the SC4 Atlas (Fig. S18f), were divided into two subpopulations in the combined atlas after projection of SC4 (Fig. 5h). An exploratory functional analysis of the differentially expressed genes in these two newly delineated megakaryocyte subpopulations (TUBA8-Mega and IGKC-Mega, Fig. S18d,e) revealed enrichment for the GO terms “humoral immune response” for IGKC-Mega cells, yet enrichment for “negative regulation of platelet activation” for TUBA8-Mega cells (Fig. 5k). These results illustrate how the continuously expandable single-cell atlases generated using SCALEX capitalize on existing large-scale data resources and also facilitate the discovery of new biological and biomedical insights.

## DISCUSSION

Single cell studies are becoming more and more prevalent, growing larger and larger in scale, and expanding in the scope of sample types, often with quite heterogenous cell subsets. Thus, there is a great need for data integration tools to accurately and efficiently handle these Atlas-level datasets^33^. Further, there is also a need for online integration capacity to continuously incorporate incoming new data with existing integrations without having to recalculate from scratch^24^. By design, SCALEX learns a generalized projection function to project heterogeneous single-cell data into a common cell-embedding space, enabling it to achieve *bona fide* online data integration. SCALEX is also computationally efficient, and preserves biological variations and avoids over-correction when integrating partially overlapping datasets.

These features make SCALEX particularly useful for Atlas-level datasets, allowing the integration of many single-cell studies to support ongoing, very large-scale research programs throughout the life sciences and biomedicine. We speculate that use of SCALEX to project single-cell datasets from highly diverse cancer types to construct a pan-cancer single-cell atlas may lead to the discovery of previously unknown cell-types that are common to divergent carcinomas and that function in pathogenesis, malignant progression, and/or metastasis.

## Methods

### Overview of the SCALEX model

SCALEX applies a variational autoencoder (VAE) to project the different batches of datasets into the same batch-invariant low-dimensional embeddings by learning a batch-free encoder and a batch-specific decoder simultaneously. Since the encoder and decoder are coupled to learn a batch-free encoder, a batch label (*b*) is only exposed to the decoder within the domain-specific batch normalization (DSBN), thus the decoder captures the batch information while the encoder learns the domain-invariant features. In the encoder, SCALEX takes the input expression profile (*x)* across all the batches as a whole mixture distribution without distinguishing their batch sources and extracts their mean (*µ*) and variance (*a^2^*) of the latent representations (*z*) in a 10-dimension embedding space to learn their global data structure. A standard multivariate Gaussian prior is used for z, while the approximated distribution of z is re-parameterized by *z = µ + a ∗ ε*, where *ε* is sampled from *ℕ(0, I)*. In the decoder, SCALEX maps the latent representations with batch label (*b*) back to their original profile. To enable the decoder to capture the batch-specific variations, a DSBN layer is applied to learn a batch-specific normalization for each batch label (*b*), before transforming them back to their original profile with the new batch variations. To learn the global distribution to avoid overcorrection on partially overlapping datasets, within each mini-batch in the training process, SCALEX randomly samples data from all batches and trains on them together with Batch Normalization to smooth the batch-specific shifts and align to the global distribution. Once trained, the encoder of SCALEX is generalized to any batches and serves as a universal function for globally mapping different batches of datasets into the same batch-invariant space.

Training SCALEX is to maximize the log-likelihood of the observed single-cell sequencing data *x*:

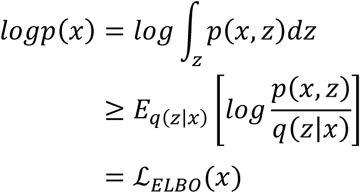

Then the loss function is transformed into the evidence lower bound (ELBO). While the ELBO can be further decomposed into two terms:

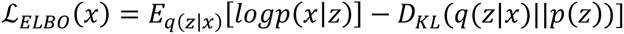

The first term is the reconstruction term, which minimizes the distance between the generated output data *x′* and the original input data *x*, calculated as the binary cross entropy between *x′* and *x*. The second term is the regularization term, which minimizes the Kullback-Leibeler divergence between posterior distribution *ℕ(µ, σ)* and prior distribution *ℕ(0, I)* of latent representations *z*. To enable a more flexible alignment under the latent space, we adjusted the coefficient of the second term to 0.5 after hyper-parameter optimization via a grid search; thus, the final loss function is:

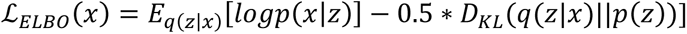

The overall network architecture of SCALEX consists of an encoder and a decoder. The encoder is a two-layer neural network (fully connected [1024]-BN-ReLU-fully connected[10]) for mean (*µ*) and variance (*a^2^*) of the 10-dimension latent representations z using a reparameterization to obtain latent representations z, and the decoder has only one layer (no hidden layer), directly connecting latent representations *z* to the output *x* (fully connected-DSBN-Sigmoid) with domain-specific batch normalization, where the latent representations z and batch ID are provided as input, and a Sigmoid activation function. We used the Adam^75^ optimizer with a 5e-4 weight decay and betas (0.9, 0.999, the exponential decay rate for the first and second moment parameters) to optimize the model under the learning rate 2e-4. We adopted mini-batch strategy to iteratively optimize the model, in each mini-batch, we randomly sampled data from all batches instead of from the same batch, and the mini-batch size for training input is 64. The maximum number of training iterations is 30,000 and an early stopping is triggered when there has been no improvement for 10 epochs. The hyper-parameters are chosen after a grid search. SCALEX is very robust with all of these hyper-parameters, all of the results in this manuscript are produced under the same parameters.

### Domain-specific batch normalization (DSBN)

Batch normalization (BN)^32^ is a widely used training technique in deep neural networks to reduce internal covariate shifting. A BN layer whitens activations within a mini-batch of samples followed by scaling and shifting with learned affine parameters γ and *β*. For a mini-batch of samples: *ℬ = {x_1…m_}*;

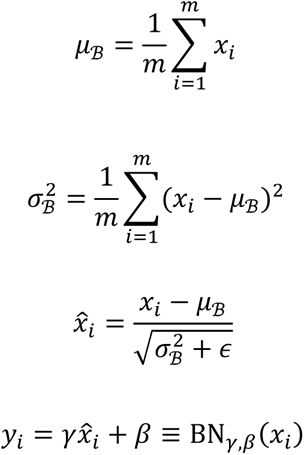

Where *µ_ℬ_*is the mini-batch mean, 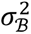 is the mini-batch variance, 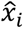 is the normalized output by *µ_ℬ_*and 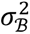, *y* is the BN output by scaling and shifting 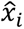 with parameters γ and β, and ε is a constant added to the mini-batch variance for numerical stability.

Domain specific batch normalization (DSBN)^31^ is a combination of multiple sets of BN specific to each domain. DSBN learns domain-specific affine parameters γ*_d_*and *β_d_*for each domain, *d* is the domain label; here, domain represents different batches. In the neural network, DSBN serves like multi-channel BN and switches to the corresponding BN given the domain label *d*. The DSBN layer can be written as:

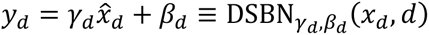

where *d* is the batch label, and γ*_d_*and *β_d_*are domain-specific affine parameters for domain *d*.

DSBN could capture the domain-specific information by estimating mini-batch statistics by learning affine parameters for each domain separately, thus enabling the network to learn the domain-invariant features.

### Preprocessing for scRNA-seq

We downloaded gene expression matrices and preprocessed them using the following procedure: i). Cells with fewer than 600 genes and genes present in fewer than 3 cells were filtered out. ii). Total counts of each cell were normalized to 10,000. iii). Values of each gene were subjected to log transformation with an offset of 1. iv). The top 2,000 highly variable genes were identified. v). Values of each gene were normalized to the range of 0-1 within each batch by the *MaxAbsScaler* function in the *scikit-learn* package in Python. The processed matrix was used as input for the SCALEX model for downstream differential gene expression analysis.

For the *human fetal atlas* dataset, we collected two batches (batch GSE156793, which contains 4,062,980 cells by sciRNA-seq3, and batch GSE134355, which contains 254,266 cells by Microwell-seq). We then selected the cells from the common tissues (1,369,619 cells) for integration and computational efficiency benchmarking (down-sampled from different data sizes including 10K, 50K, 250K, 1M, and 4M).

### Preprocessing for scATAC-seq

We downloaded open chromatin profile matrices (peaks or bins), merged them by peaks (or bins), and processed them using the following procedure: i). The combined matrix was binarized and filter bins with fewer than 3 cells. ii). The top 30,000 most variable peaks (or bins) were selected using the *select_var_feature* function in the *EpiScanpy*^76^ package. iii). Total counts of each cell were normalized to the median of the total counts of all cells by using the *normalize_total* function, with parameters *target_sum*=“None” in the *Scanpy*^77^ package. iv). Values of each peak (or bin) were normalized to the range of 0-1 within each batch by the *MaxAbsScaler* function in the *scikit-learn* package in Python. The processed matrix was used as input for the SCALEX model.

### Preprocessing for cross-modality data (scRNA-seq and scATAC-seq)

We first created a gene activity matrix by the *GeneActivity* function in the *Signac*^78^ R package to quantify the activity of each gene from scATAC-seq data. We then combined gene activity score matrix with scRNA-seq data matrix as two individual “batches” for integration. The subsequent preprocessing followed the same preprocessing used for the scRNA-seq data (above).

### Visualization

UMAP algorithm^46^ was used for visualization. We applied the *neighbors* function from the Python package *Scanpy* with the parameters *n_neighbors*=30 and *metric*=“Euclidean” for computing the neighbor graph, followed by *umap* function with *min_dist*=0.1 to visualize cells in a two-dimensional space.

### Adjusted Rand Index

The Rand Index (RI) computes a similarity score between two clustering assignments by considering matched and unmatched assignment pairs, independent of the number of clusters. The Adjusted Rand Index (ARI) score is calculated by “adjust for chance” with RI as follows:

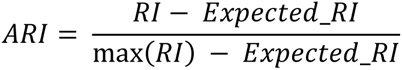

If given the contingency table, then ARI can also be represented by:

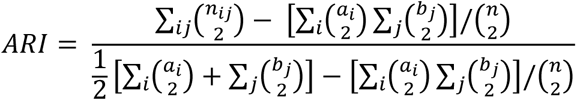

The ARI score is 0 for random prediction and 1 for perfectly matching.

#### Normalized mutual information

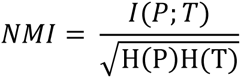

Where P and T are categorical distributions for the predicted and real clustering, I is the mutual entropy, and H is the Shannon entropy.

### Silhouette score

We used the silhouette score to assess the separation of biological populations with the function *silhouette_score* in the *scikit*-*learn* package in Python. The silhouette score was computed by combining the average intra-cluster distance (a) and the average nearest-cluster (b) for each cell.

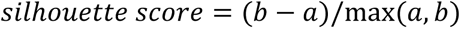

Here, we took UMAP embeddings as input to calculate silhouette score.

### Batch entropy mixing score

Batch entropy mixing score (adapted from “entropy of batch mixing”^19^) was used to access the regional mixing of cells from different batches, with a high score suggesting that cells from different batches are well mixed together.

The batch entropy mixing score was computed as follows:

1. Calculated the proportion *Pi* of cell numbers in each batch to the total cell numbers.
2. Randomly chose 30 cells from all batches.
3. Calculated the 30 nearest neighbors for each randomly chosen cell.
4. The regional mixing entropies for each cell were defined as:

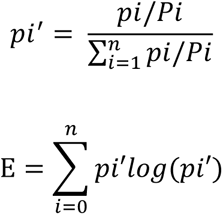

where *pi* is the proportion of cells from batch *i* in a given region, such that 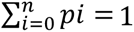, *pi*’ is a correction item to eliminate the deviation caused by the different cell numbers in different batches. The total mixing entropy was then calculated as the sum of the regional mixing entropies.
5. Repeated (2)-(4) for 10 iterations with different randomly chosen cells and calculated the average, E, as the final batch entropy mixing score.

Note that to mitigate the effect of misalignment of batch-specific cell-types, we calculated the batch entropy mixing score only based on cells from cell-types that are common in different batches.

### Local inverse Simpsons Index (LISI)

The LISI metric was proposed by Korsunsky *et al. 2019* ^20^ to assess batch and cell-type mixing. We calculated integration LISI (iLISI) and cell-type LISI (cLISI) values using the *compute_lisi* function in the *lisi* R package. UMAP embeddings, batch labels, and cell-type labels were used as input in calculation. Briefly,

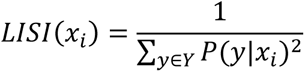

where *x_i_∈ {x_1_, x_2_,…, x_N_}*is the *i*-th cell’s UMAP embeddings in the dataset of size *N*, and *Y* is the set of unique values with respect to the type of LISI we are computing (*i.e.*, *Y* is the values of “batch label” for calculating iLISI and the value of “cell-type label” for calculating cLISI). The probability *P(y|x_i_)*refers to the “relative abundance” of the covariate *y* within KNN (k-nearest neighborhood) of *x_i_*. A Gaussian kernel-based distribution of neighborhoods was used and the perplexity was fixed to 30.

### Over-correction score

We defined an over-correction score to assess the level of over-correction problem, based on calculating the percentage of cells with inconsistent cell-types in each cell’s neighborhood. We calculated the over-correction score over all cells, and for each cell i we averaged the frequency of the k-nearest neighboring cells with distinct cell-types to the cell i (see the following equation).

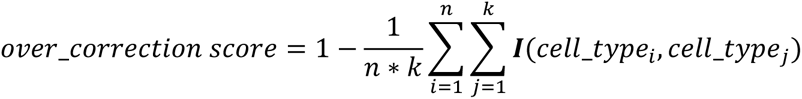

where n is the total cell number, k represents the k-nearest neighbors of each cell, the cell-type of the cell i is *cell_type_i_*, the cell-type of the neighboring cell j is *cell_type_j_*, and *I* is an indicative function defined as:

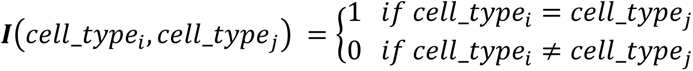

Formally, the over-correction score is a negative index, i.e., the higher the over-correction score, the more severe the extent of inaccurate mixing of cell-types.

### Comparison with other integration methods

We compared SCALEX to nine other batch effect removal methods (see below for specific details of each method). For each dataset as input for all methods, we performed the same filtration, followed by method-specific normalization, batch correction and visualization. Note that for visual comparison, we also included the embeddings of the raw input data, wherein we performed dimensionality reduction by Principal Component Analysis (PCA)^79^ followed by UMAP visualization to see the batch effects. No correction function was used. All parameters were kept as default values.

Scanorama (v1.6). We performed the preprocessing pipelines as stated above (as the same below), and used the *Scanpy* and *scanorama* Python packages for integration. For the *highly_variable_genes* function, we set *flavor*=“seurat”, *batch_key*=“batch”, and *n_top_genes*=2,000. After extracting highly variable genes, we divided the datasets according to the batch labels and formed a new list of datasets as the input for the *correct_scanpy* function. The integration matrix was kept for downstream analysis. All other parameters were kept their default values.

BBKNN (v1.3.12). We used *Scanpy* and *bbknn* Python packages and followed the suggested pipelines for integration. For the *highly_variable_genes* function, we set *flavor*=“seurat”, *batch_key*=“batch”, and *n_top_genes*=2,000. After selecting cell neighbors at the low-dimensional space from the PCA analysis, we performed the *bbknn* function with *neighbors_within_batch*=5, *n_pcs*=20, and *trim*=0. All other parameters were default.

scVI (scvi-tools, v0.11): We used the *scvi* Python package and followed the suggested pipelines. Batch information was added to the VAE model by setting n_batch.

Seurat v3 (v3.2.3): We used the *Seurat* R package and followed the standard integration workflow. We normalized different batches of a dataset separately. For the *FindVariableFeatures* function, we set *selection.method*=“vst” and *nfeatures*=2,000 to select 2,000 highly variable genes for each batch of a dataset. For the *FindIntegrationAnchors* function, we set *k.filter*=100. All other parameters were kept at default values. If the number of input cells in a dataset exceeded 50,000, we employed the reciprocal PCA and reference-based integration to improve computational efficiency.

Harmony (v1.0): We used the *harmony* R package. We created a *Seurat* object with all cells and performed the standard workflow. After PCA, we used the *RunHarmony* function for integration. All parameters were default.

Conos (v1.3.1): We used the *Conos* R package. For each batch of dataset, we used the *basicSeuratProc* and *RunTSNE* functions for preprocessing. After that, we built a joint graph using the *buildGraph* function with *k*=30 and *k.self*=5. All other parameters were default.

MNN (FastMNN, v0.3.0): We used the *SeuratWrappers* R package. We created a Seurat object with all cells and performed the standard workflow. Then we used the *RunFastMNN* function with default parameters for integration.

Online iNMF and LIGER (LIGER, v1.0.0): We used the *rliger* R package. For the online iNMF method, we used the online_iNMF function with *k*=20, *miniBatch_size*=5,000 and *max.epochs*=5. For the LIGER method, we used the *optimizeALS* function with *k*=20. All other parameters were the default values. Different from other methods, online iNMF only loads one mini-batch from the whole data in the HDF5 file format (converted from the orginal data format by the *rhdf5* R package) for a memory-efficient implementation; accordingly, a file conversion issue with the down-sampled *human fetal atlas* dataset of 4M data size prevented online iNMF from calculating computational efficiency with the 4M.

### Cell-type annotation by clustering

This type of annotation was used for *de novo* annotation of a single-cell dataset. We used a Leiden clustering^80^ method for cell clustering (specifically employing the *leiden* function from the Python package *Scanpy* with default parameters). Then for each cluster, we annotate its cell-type based on: i) cell-type annotations of each cell in the original study, if available, or ii) expression levels of canonical marker genes in each cell. A majority vote strategy was used when needed. Similar to Ren et al. 2021, we also employed a hierarchical annotation strategy, *i.e.*, we first clustered all cells in a dataset into several major clusters, then for some big clusters, we further clustered them into minor clusters respectively.

### Single cell projection

We defined single cell projection as the operation to convert high-dimensional single-cell data (*e.g.*, gene expression profiles in scRNA-seq or open chromatin profiles in scATAC-seq) to low-dimensional representations in the common SCALEX cell-embedding space using the trained encoder.

### Cell-type annotation by label transfer

This type of annotation was used for annotation of a new single-cell data batch using the annotations in a large single-cell dataset as a reference, or for *post hoc* annotations of unknown cell population(s) in a large dataset using new batches of data of known cell-types. Both scenarios require “single cell projection” (see details below). The basic idea of cell-type annotation by label transfer is based-on that the same cell-types will occupy the same locations in the low-dimensional SCALEX cell-embedding space, thus cell-type annotation in one data batch can be transferred to another data batch, for the cells positioned at the same locations. Technically, we used the *KNeighborsClassifier* function from the *scikit-learn* package to train a prediction model, using the representations (in the low-dimensional cell-embedding space) of the single-cell data with known cell-type labels as input. We then used this model to make cell-type predictions for cells without annotations using their representations (in the low-dimensional cell-embedding space) as input. For comparison, label transfer for online iNMF follows the same procedures as SCALEX by predicting the cell-type based-on the projected locations.

### Similarity matrix and confusion matrix

We used similarity matrix to evaluate the congruence of two different batches for the same cell-types in the common cell-embedding space. Technically, we merged all cells with the same cell-type label and calculated an average representation (in the low-dimensional cell-embedding space) for the cell-type. This was repeated for all cell-types. We then calculated the similarity matrix *S=[S_ij_]* for the cell-type similarities of the two batches, where *S_ij_* is the Pearson correlation coefficient between the average representation of cell-type *i* in data_batch_1 and the average representation of cell-type *j* in data_batch_2.

We used the confusion matrix to evaluate the accuracy of cell-type annotations (prediction) when a gold-standard annotation is available, which is typical for “cell-type annotation by label transfer” (see above). In cell-type annotation by label transfer, we predict the cell-types for a single-cell data_batch_1, using the annotations in another data_batch_2. When data_batch_1 was already annotated with cell-types, we can calculate the confusion matrix *C=[C_ij_]* to compare the cell-type predictions with the existing cell-type annotations, where *C_ij_* equals the percentage of cells known to be in cell-type *i* and predicted to be in cell-type *j*.

### F1 score

F1 score is a weighted average of the precision and recall as a classification metric, which is computed by:

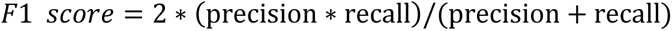

### Generation of partially overlapping datasets

To simulate partially overlapping datasets from the *pancreas* dataset, we used the pancreas_celseq2 and pancreas_smartseq2 data batches, and worked with only six cell-types (alpha, beta, ductal, acinar, delta, gamma). For each simulated partially overlapping dataset, we randomly selected three to six cell-types from each batch, and counted the number of the common cell-types, which was used as the indicator for the overlapping level (whole integers, 0 to 6). We required the union of cell-types in the newly simulated partially overlapping dataset to cover all six cell-types.

For the *PBMC* dataset, we used both of the two data batches and worked with twelve cell-types (B, CD4 T, CD4 naive T, CD8 T, CD8 naive T, DC, HSC, Megakaryocyte, NK, monocyte-CD14, monocyte-FCGR3A, pDC). We used the same down-sampling strategy as for the *pancreas* dataset (above).

### Analysis of changes in cell-type frequency across multiple conditions

To identify differences in cell-type frequency among the scRNA-seq data from the mild/moderate, severe, convalescent COVID-19 patients, as well as the healthy and influenza patient controls, we applied a Dirichlet-multinomial regression model. This model accounts for the constraint that the cell frequencies in a scRNA-seq data are not independent of each other. In detail, we normalized the regression coefficients to a standard normal distribution and calculated a z score, and then conducted significance testing based on the regression model generated by the *DirichReg* function in the R package *DirichletReg* (v 0.7).

### Differential gene expression analysis and Gene Ontology term enrichment analysis

Differential gene expression analysis was performed on all expressed genes using the *rank_genes_groups* function with *method*=“t-test” in the *Scanpy* package, for two certain cell-types in a COVID-19 single-cell atlas. A gene was considered differentially expressed when a log2-fold change was >1 in the two conditions in comparison, and the Benjamini-Hochberg adjusted P-value was < 0.01. The top 200 highly expressed genes sorted by scores (implemented in *Scanpy*) of each cell-type were used as the input for GO analysis, and enriched GO terms were acquired for each group of cells of the “GO_Biological_Process_2018” dataset using the Python package *GSEApy*.

### Inflammatory and cytokine score analysis

We defined the inflammatory score and the cytokine score for each cell following Ren et al. 2021, based on the expression of a defined collection of cytokine genes and inflammatory-response-related genes, which were from Liberzon et al. 2015 (Supplementary Table 1). We then calculated the cytokine and inflammatory scores from the raw gene expression profile using the *score_genes* function implemented in the *Scanpy*.

### Ablation studies using test-variants of SCLAEX

We evaluate the contributions of each design element by the performance of different ablation variant models on integration and projection tasks (Fig. S18, Fig. S19).

**Baseline/full SCLAEX.** The baseline or the full SCALEX model is described in the paper, including an encoder without batch label, mini-batch sampling from all batches with Batch Normalization, a decoder with DSBN, and beta=0.5. All other variant models are compared to this baseline model.

**A variant of encoder with batch label.** The baseline model with an encoder with batch label as input.

**A variant of sampling by batch without Batch Normalization.** The baseline model without the Batch Normalization layer in the encoder as well as each mini-batch is sampled by batch instead of from all batches.

**A variant of decoder without DSBN.** The baseline model without the DSBN layer in the decoder.

### Software availability

SCALEX is available at https://github.com/jsxlei/SCALEX.

### Data availability

All data analyzed in this study are publicly available; the data sources are detailed in Supplementary Table 2.

## Acknowledgements

We thank Jianbin Wang, Jin Gu and Fuchou Tang for helpful comments and advice. This work is supported by the State Key Research Development Program of China (Grant No. 2018YFA0107603 and 2019YFA0110002); the National Natural Science Foundation of China (Grants No. 91740204, 91940306, and 31761163007); the Beijing Advanced Innovation Center for Structural Biology; and the Tsinghua-Peking Joint Center for Life Sciences.

## Author contributions

Q.C.Z. conceived and supervised the project. L.X. designed and implemented the SCALEX model. L.X. and K.T. validated the SCALEX model. L.X., K.T., and Y.L. analyzed the results. L.X. and Q.C.Z. wrote the manuscript, with inputs from all the authors.

## Supplementary figures

**Fig. S1.**
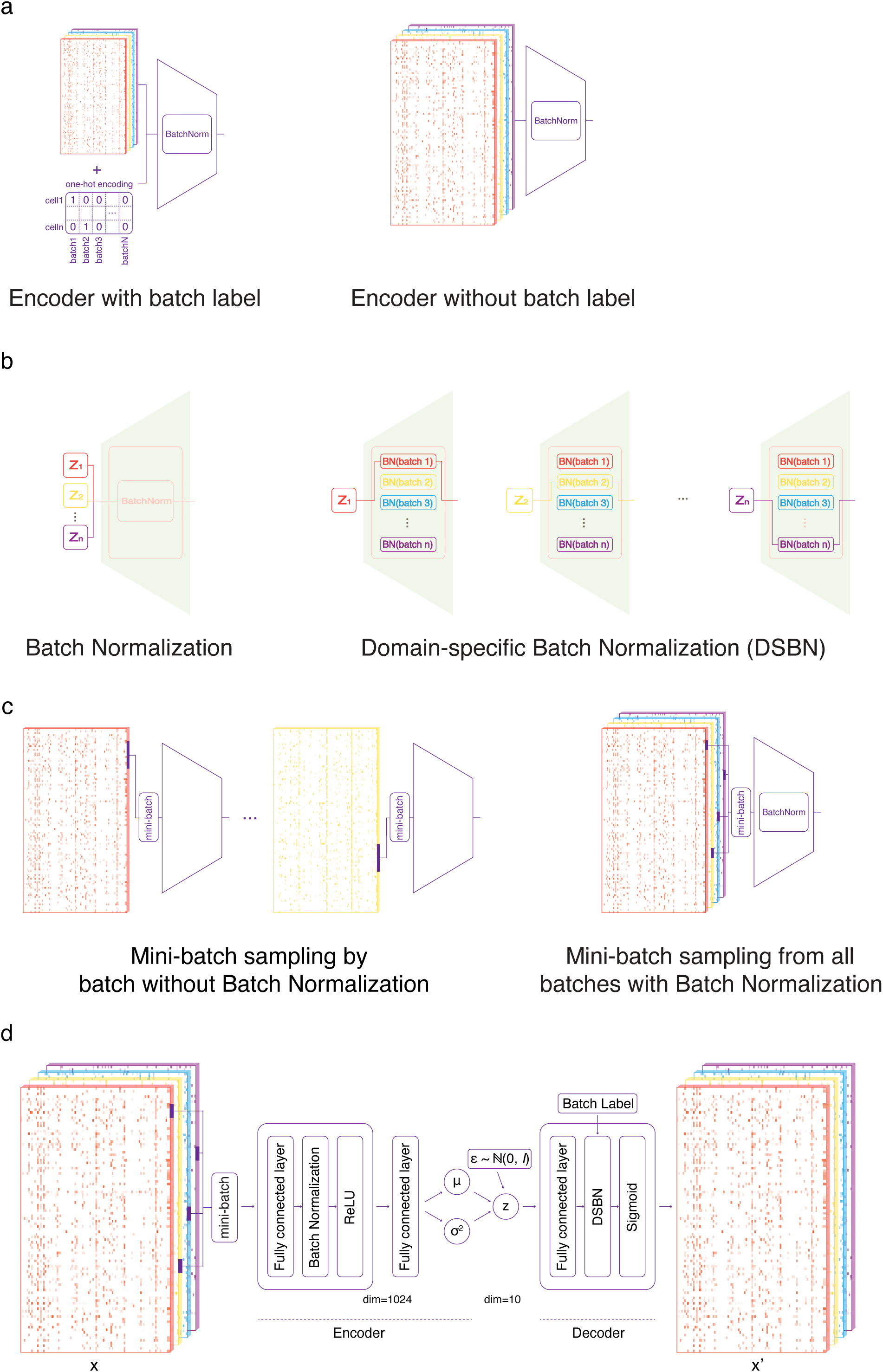
Three design elements to learn a generalized encoder. **a**, Assessing encoders with or without batch labels. **b**, Assessing decoders with or without DSBN. **c**, Assessing mini-batch sampling by batch without Batch Normalization or sampling from all batches with Batch Normalization. **d**, Detailed architecture of SCALEX and information flow.

**Fig. S2.**
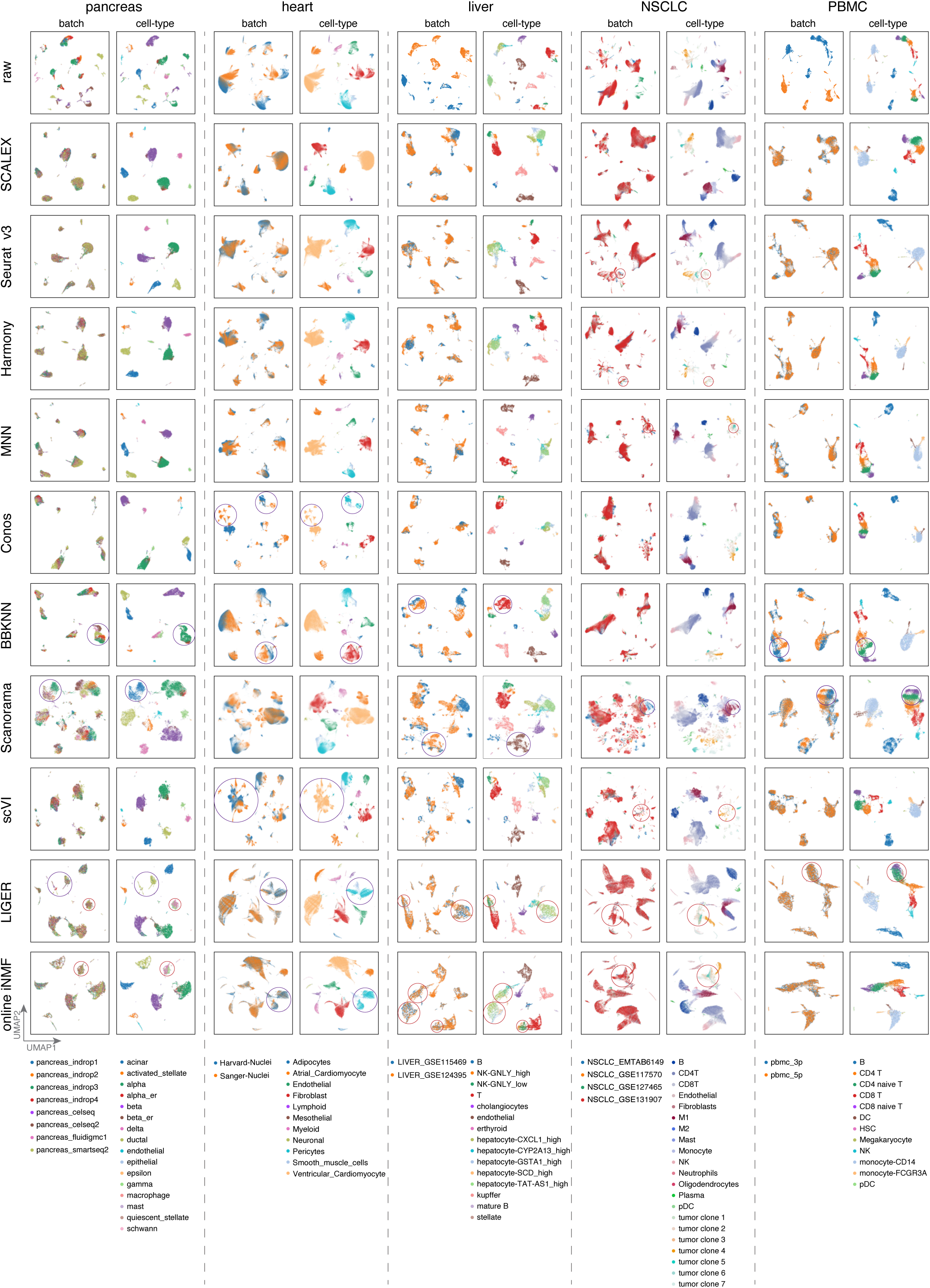
Comparisons of integration performance of indicated methods across the indicated benchmark datasets. UMAP embeddings of the indicated methods across the indicated benchmark datasets colored by batches and cell-type. Misalignments are highlighted with red circles.

**Fig. S3.**
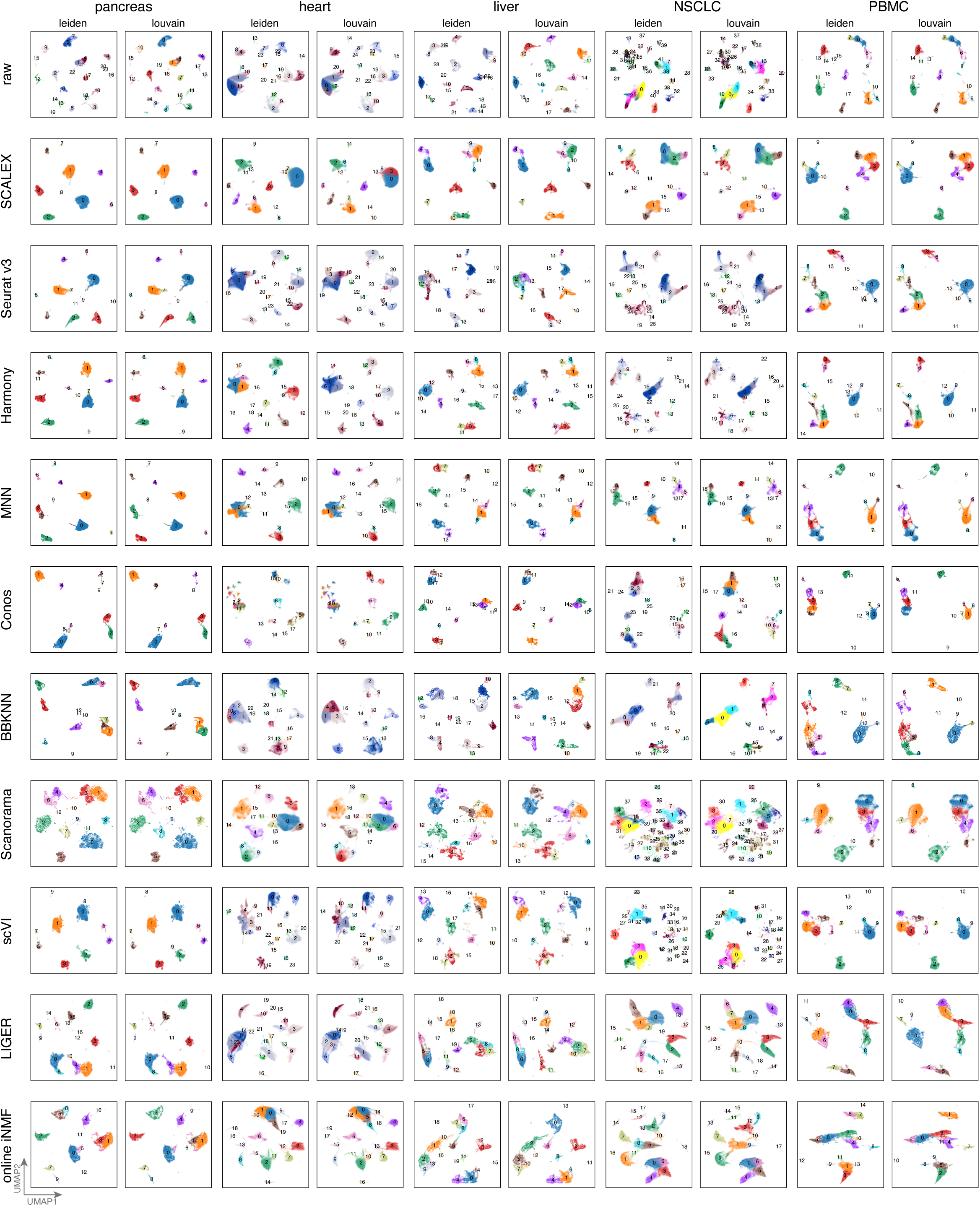
Comparisons of clustering results of indicated methods across the indicated benchmark datasets. UMAP embeddings of clustering results of the indicated benchmark datasets after integration by indicated methods; colored by Leiden (left) and Louvain (right).

**Fig. S4.**
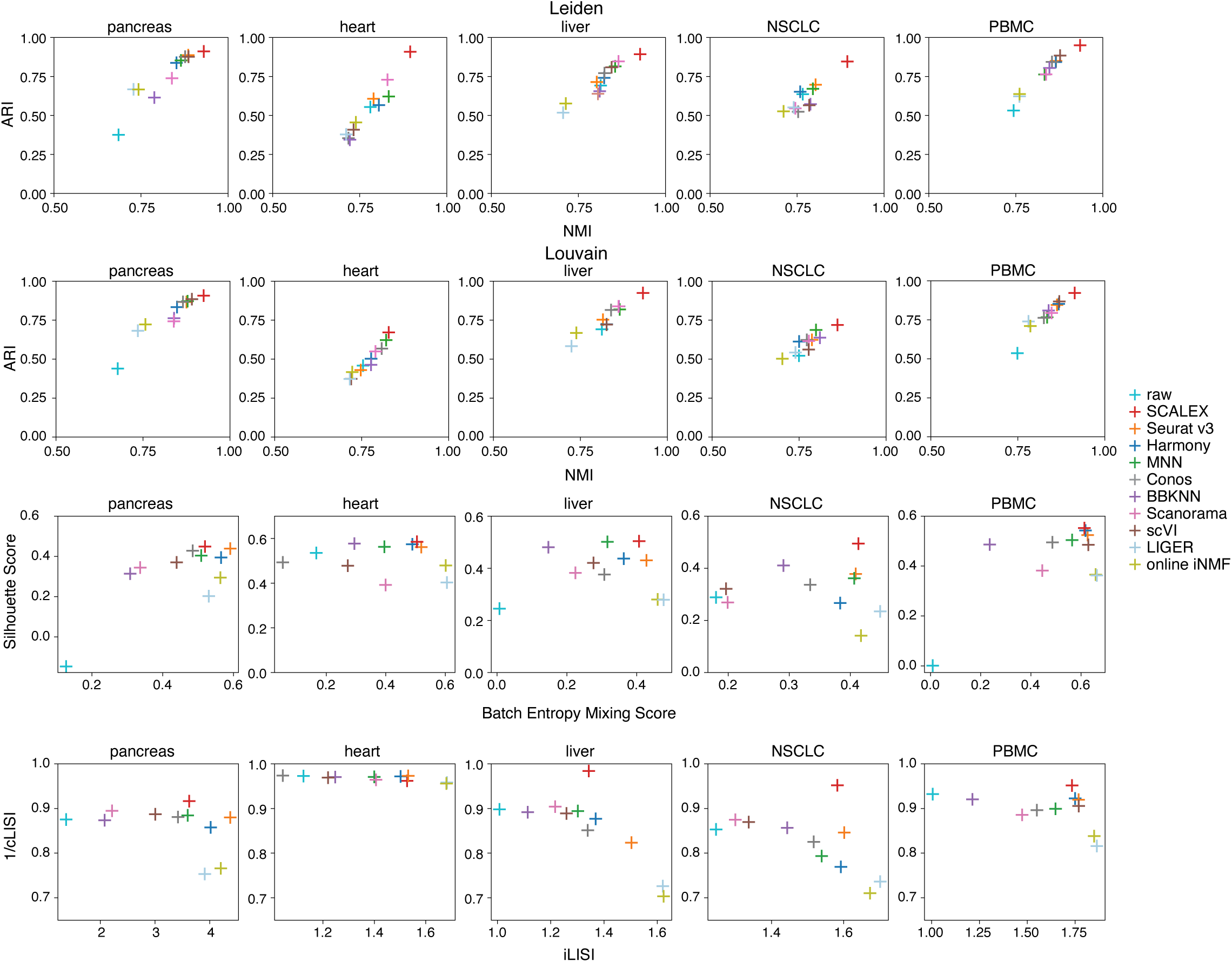
Comparisons of integration performance by quantification metrics. Scatter plots showing the ARI and NMI scores based-on the Leiden and Louvain clustering results, the Silhouette and batch entropy mixing scores, and the cLISI/ iLISI scores.

**Fig. S5.**
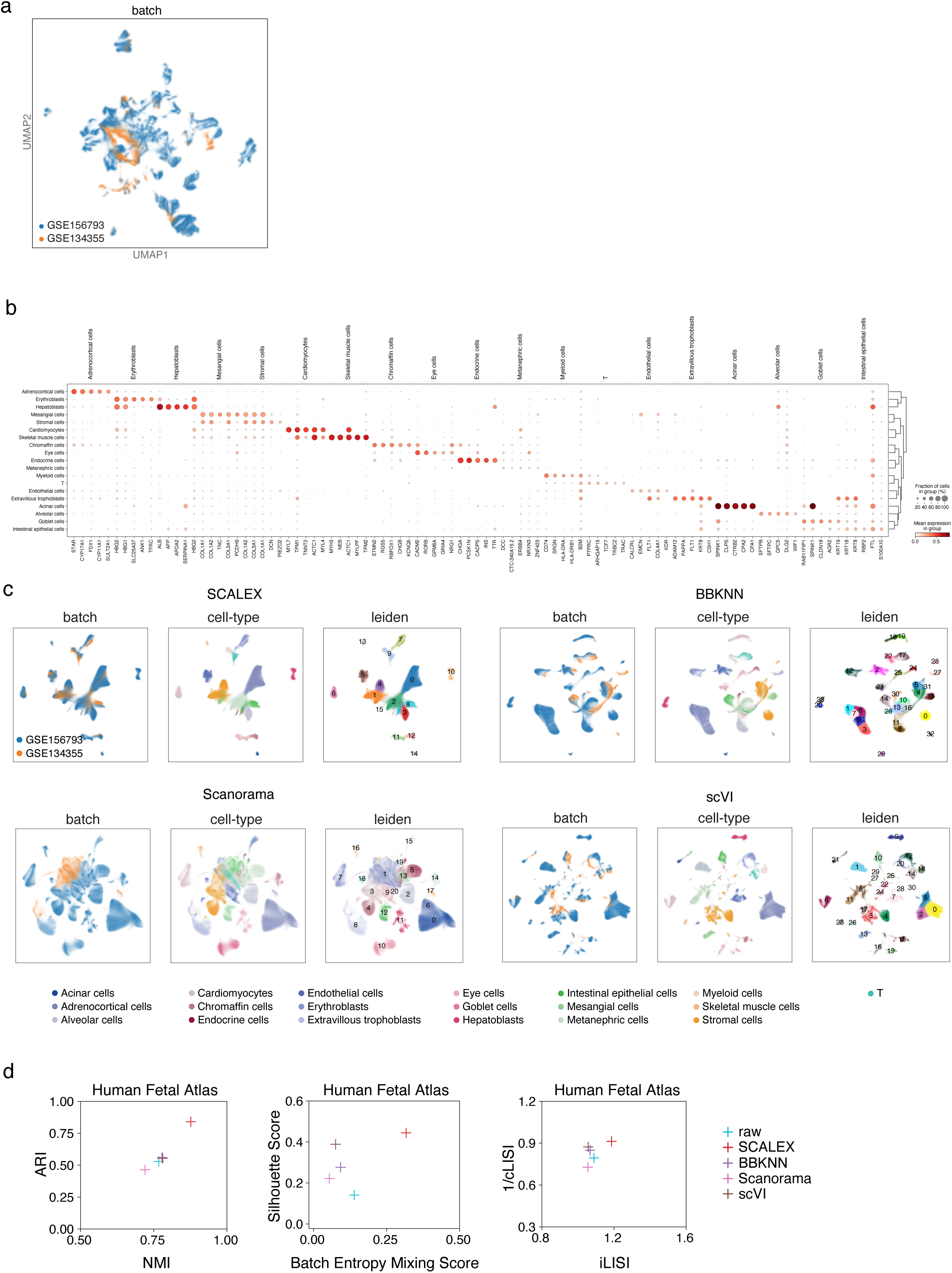
Comparisons of integration performance of indicated methods based on Human Fetal Atlas dataset. **a**, UMAP embeddings of the Human Fetal Atlas before integration. **b,** Dotplot of canonical marker genes for each cell-type. Dot color represents average expression level, and dot size represents the proportion of cells in the group expressing the marker. **c**, UMAP embeddings of the indicated methods (only SCALEX, BBKNN, Scanorama, and scVI are scalable to the Human Fetal Atlas dataset). Cells are colored by batch (left), cell-type (middle), and Leiden clustering (right). **d**, Scatter plot showing the ARI and NMI scores based-on the Leiden clustering results, the Silhouette and batch entropy mixing scores, and the iLISI and 1/cLISI scores. Note that online iNMF was not successfully tested on the Human Fetal Atlas due to a HDF5 file conversion issue for large data.

**Fig. S6.**
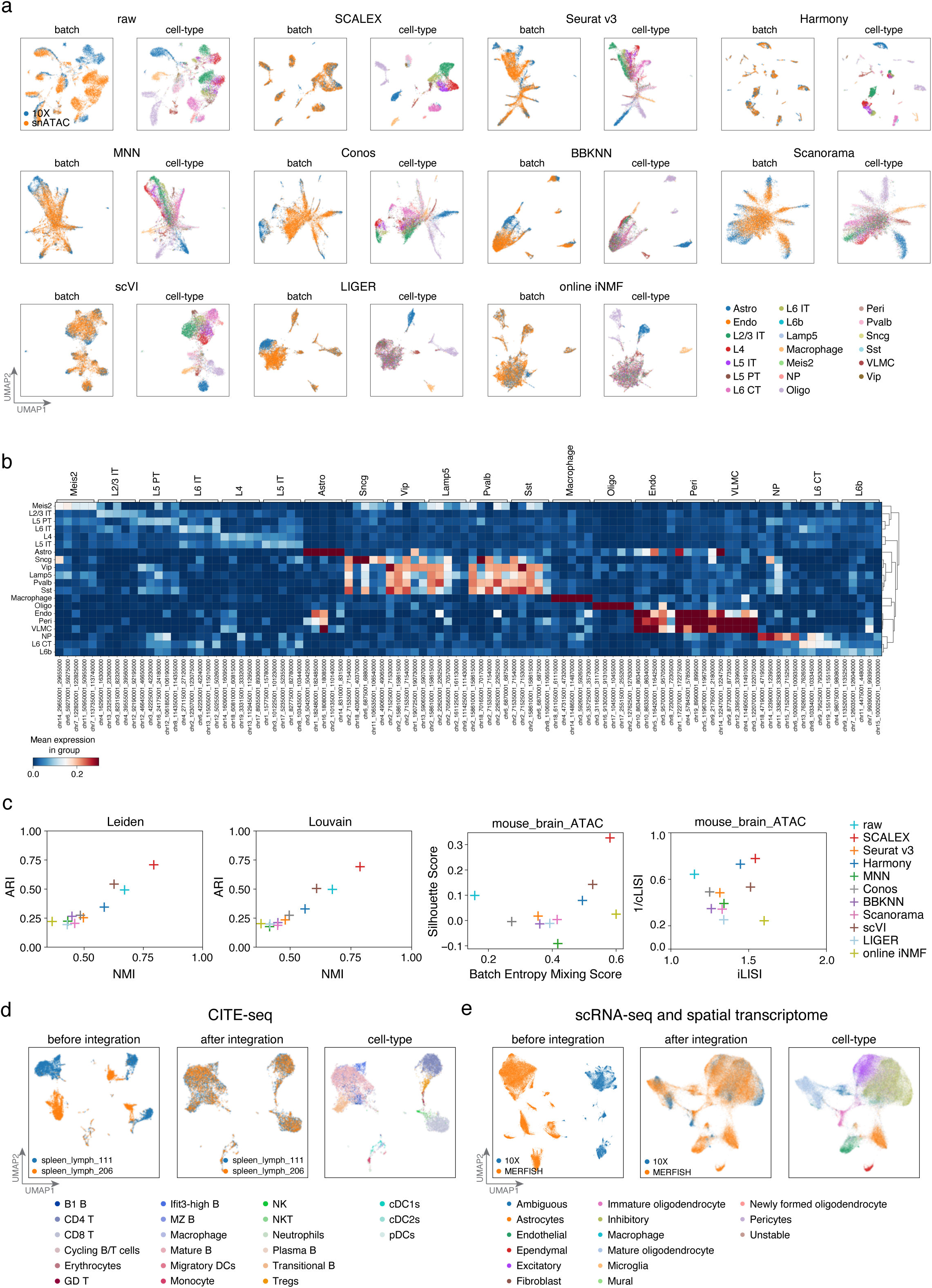
Comparisons of integration performance of indicated methods based on scATAC-seq dataset and other modality datasets. **a**, UMAP embeddings of the mouse brain scATAC-seq dataset before and after integration by indicated methods; colored by batch (left) and cell-type (right). **b**, Cell-type-specific peaks for the mouse brain scATAC-seq dataset. **c**, Scatter plot showing the ARI and NMI scores based-on the Leiden and Louvain clustering results, the Silhouette and batch entropy mixing scores, and the iLISI and 1/cLISI scores. **d**,**e**, UMAP embedding before and after SCALEX integration over CITE-seq (which measures abundance of both RNA and protein in single cells) spleen lymph dataset and cross-modality mouse brain dataset between spatial transcriptome (MERFISH) dataset and RNA-seq (10X) dataset.

**Fig. S7.**
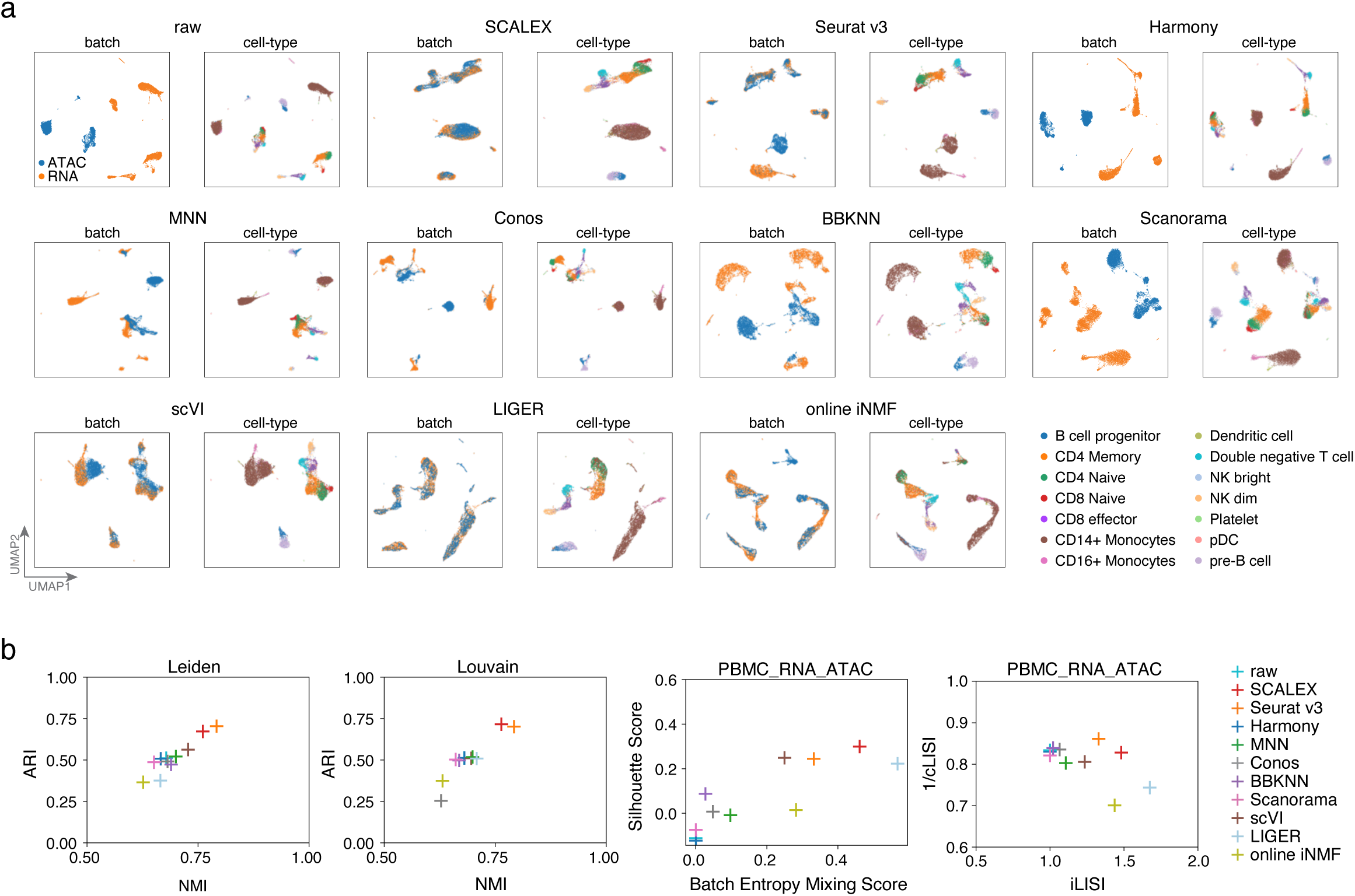
Comparisons of integration performance of indicated methods based on cross-modality dataset. **a**, UMAP embeddings of the PBMC cross-modality dataset by indicated methods. Cells are colored by batch or cell-type. **b**, Scatter plot showing a quantitative comparison of the silhouette score (y-axis) and the batch entropy mixing score (x-axis) (left), ARI (y-axis), and NMI (x-axis) based-on Leiden clustering (middle) and Louvain clustering (right), the Silhouette and batch entropy mixing scores, and the iLISI and 1/cLISI scores for the PBMC cross-modality dataset.

**Fig. S8.**
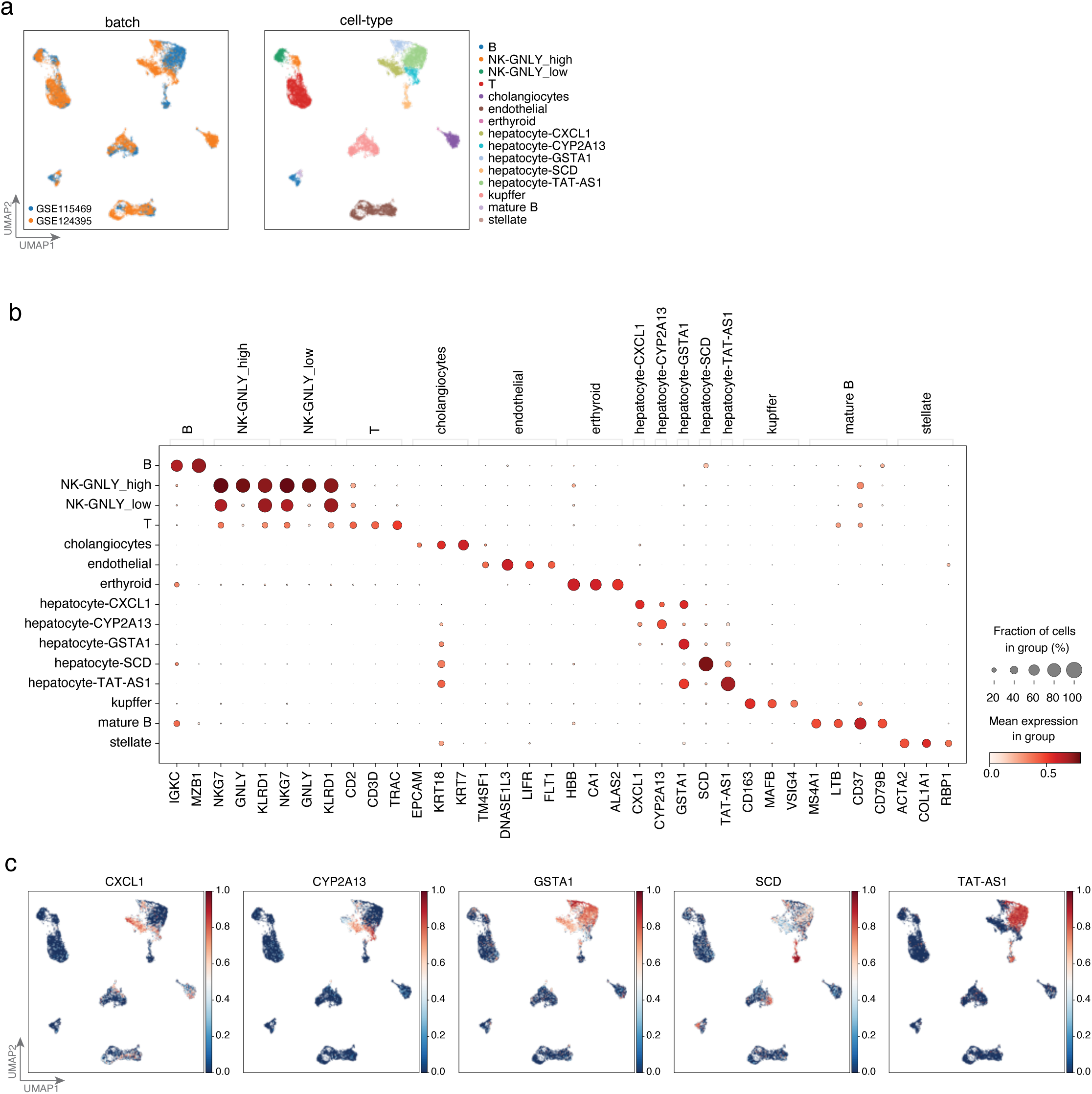
Canonical marker genes of different cell-types and UMAP embeddings of the *liver* dataset. **a,** UMAP embeddings of the *liver* dataset, colored by batch (left) and cell-type (right) after SCALEX integration. **b,** Dotplot of canonical marker genes for each cell-type. Dot color represents average expression level, and dot size represents the proportion of cells in the group expressing the marker. **c**, Normalized marker gene expression on the UMAP embeddings of the five hepatocyte subtypes. Color bar represents the expression level.

**Fig. S9.**
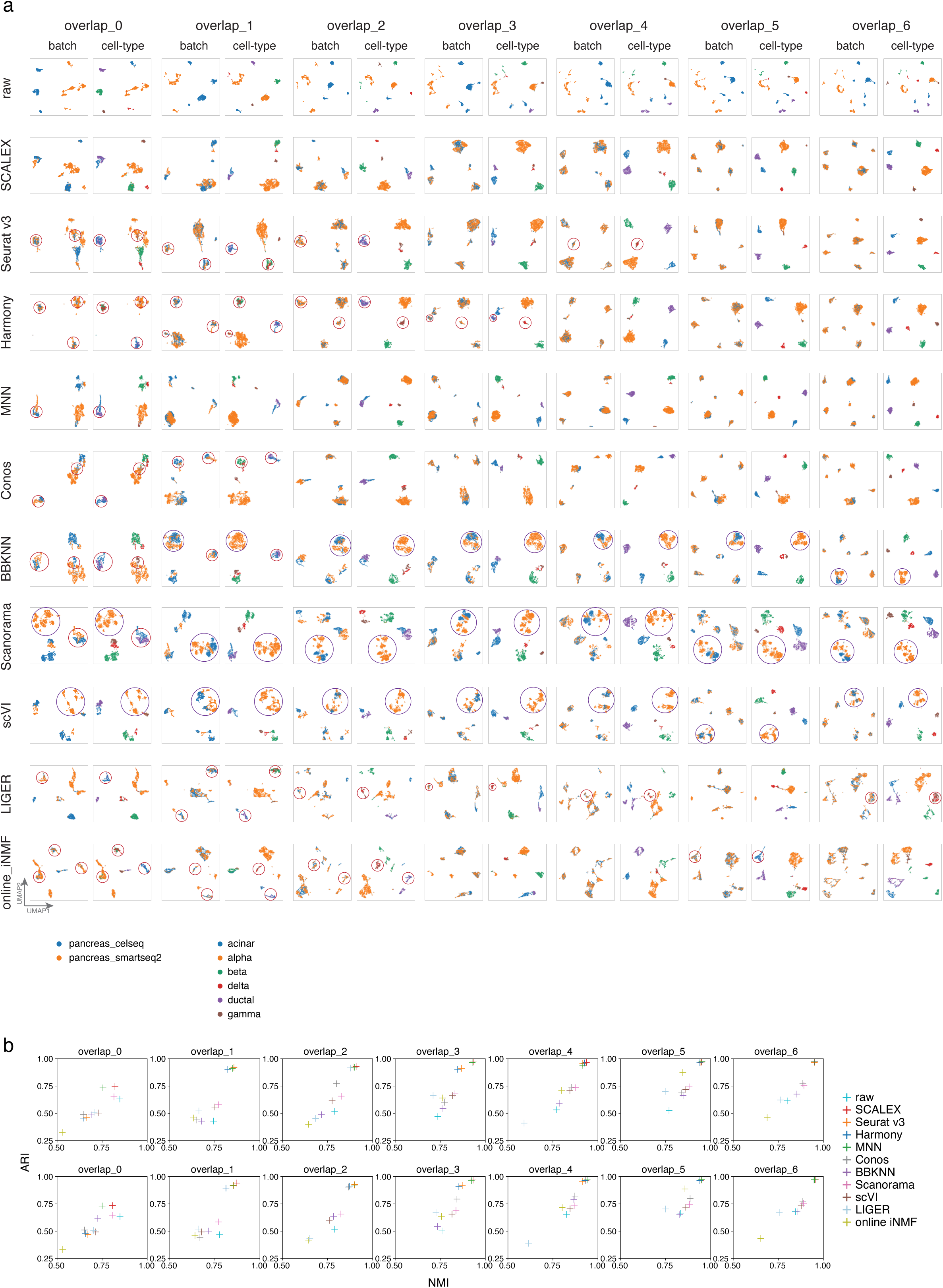
Comparisons of integration performance based on partially overlapping simulated *pancreas* dataset. **a,** Partially overlapping datasets were generated by down-sampling the *pancreas* dataset, consisted of common cell-types with a decreased overlapping number (ranging from 0 to 6). Integration results for SCALEX, Seurat, Harmony, and online iNMF are shown in the UMAP embeddings colored by batches (left) and cell-types (right) respectively (overlapping number decreases from 6 to 0). Misalignments are highlighted with red circles. **b**, Scatter plot showing a quantitative comparison of the indicated methods in terms of the ARI score (y-axis) and the NMI score (x-axis), based on the Leiden (top) and Louvain (bottom) clustering results in the latent space based on simulated *pancreas* datasets.

**Fig. S10.**
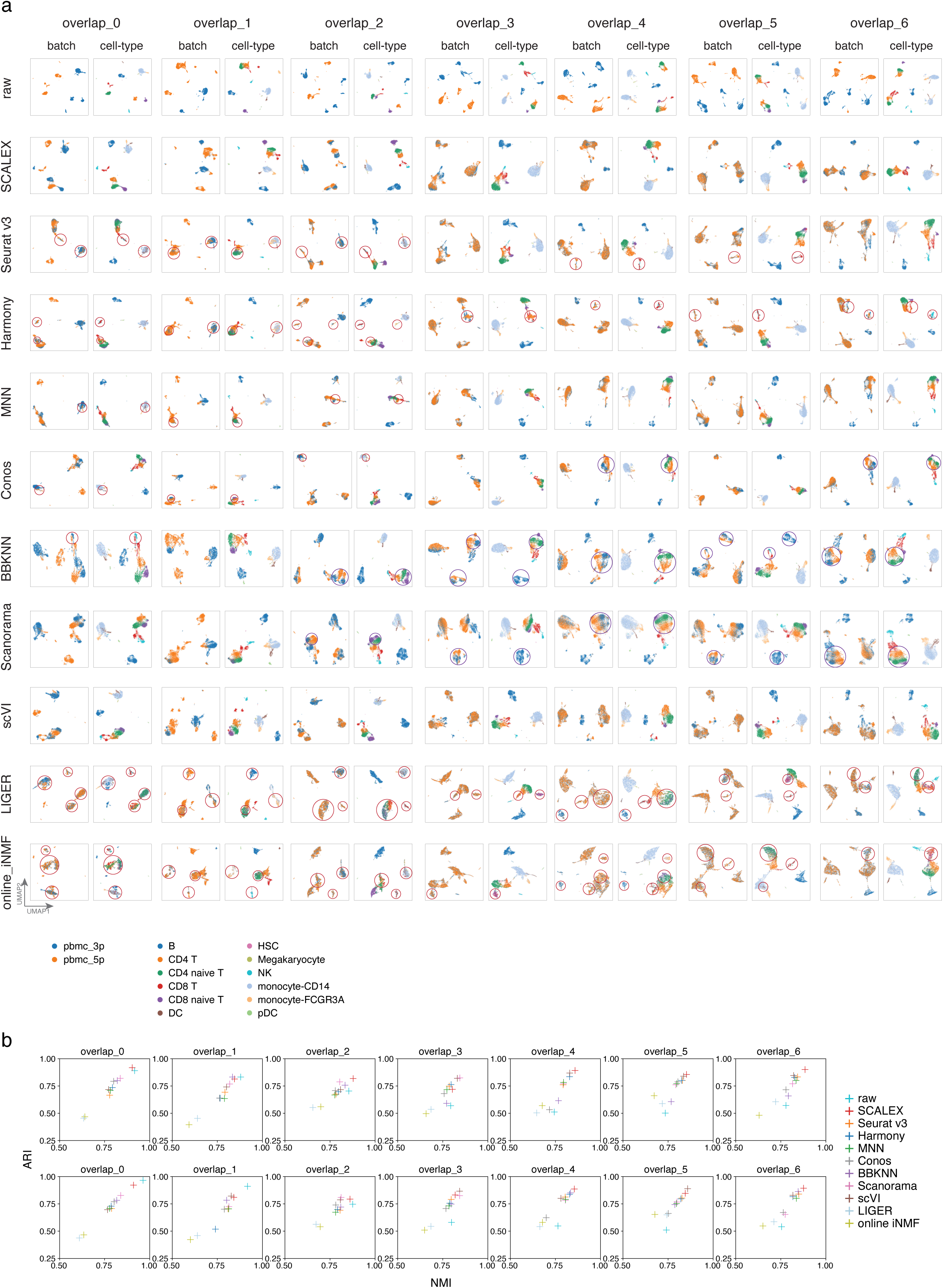
Comparisons of integration performance based on partially overlapping simulated *PBMC* dataset. **a**, Partially overlapping datasets were generated by down-sampling the *PBMC* dataset, consisted of common cell-types with a decreased overlapping number (ranging from 0 to 6). Integration results for SCALEX, Seurat, Harmony, and online iNMF are shown in the UMAP embeddings colored by batches (left) and cell-types (right) respectively (overlapping number decreases from 6 to 0). Misalignments are highlighted with red circles. **b**, Scatter plot showing a quantitative comparison of the indicated methods in terms of the ARI score (y-axis) and the NMI score (x-axis), based on the Leiden (top) and Louvain (bottom) clustering results in the latent space based on simulated *PBMC* datasets.

**Fig. S11.**
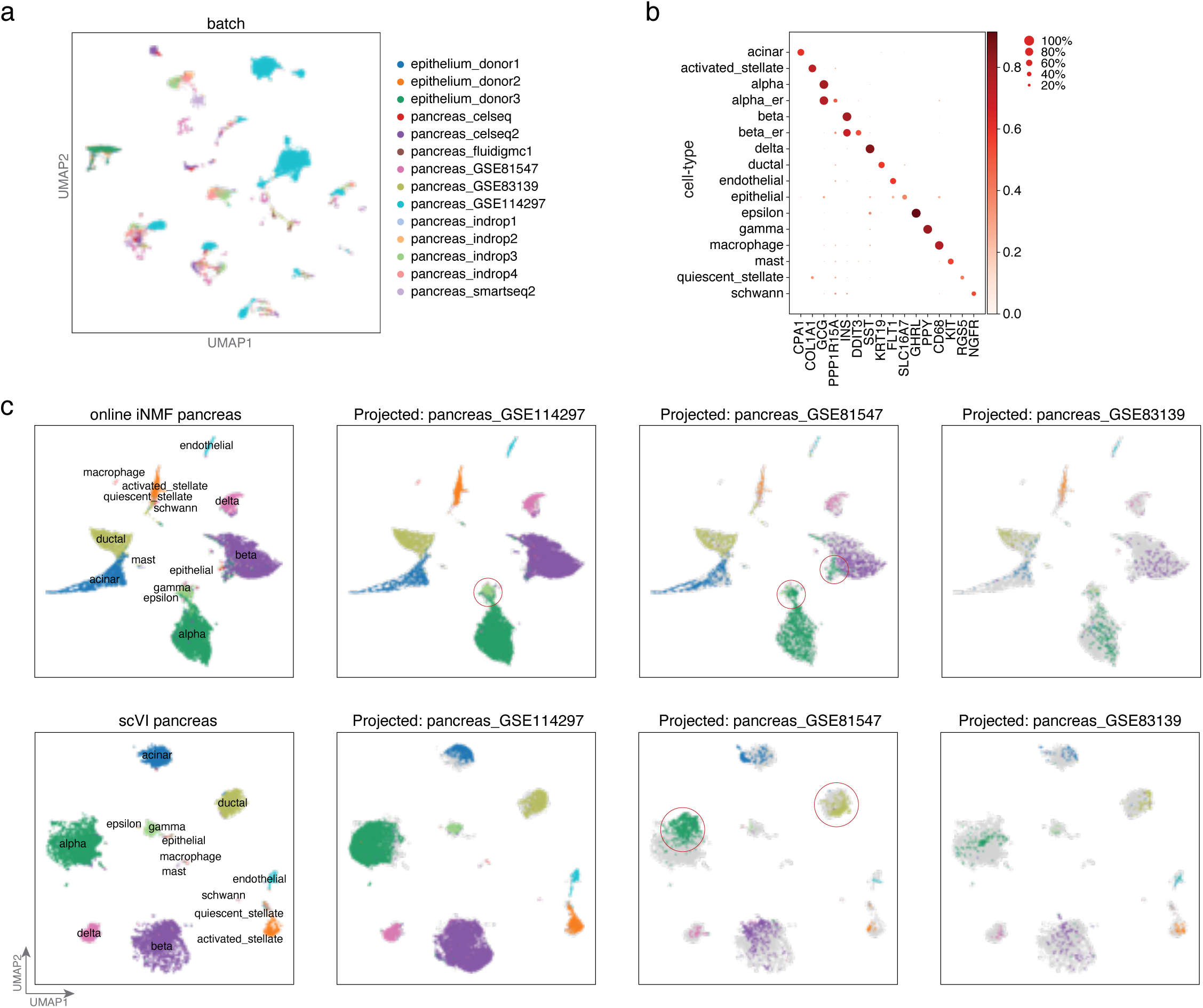
Projection of three additional pancreas data batches onto the *pancreas* dataset. **a**, UMAP embeddings of the *pancreas* dataset and the three additional pancreas data batches and the bronchial epithelium data batches (data from three donors) before integration. Cells are colored by batch. **b**, Dot plot of canonical markers of cell-types of reference *pancreas* dataset; dot color represents average expression level, and dot size represents the proportion of cells in the group expressing the marker. **c,** UMAP embeddings of the common cell space obtained by using online iNMF (top) and scVI (bottom) to project the three additional indicated pancreas data batches onto the *pancreas* dataset. Cells are colored by cell-type with light gray shadows representing the original *pancreas* dataset. Misalignments are highlighted with red circles. Note that here for convenient comparison of the projected data with the existing data, we showed the UMAP embeddings of the *pancreas* dataset combined with the projected data, which are visualized differently from the UMAP embeddings of the *pancreas* dataset alone in Fig. S2 as UMAP visualizations change with embedding data.

**Fig. S12.**
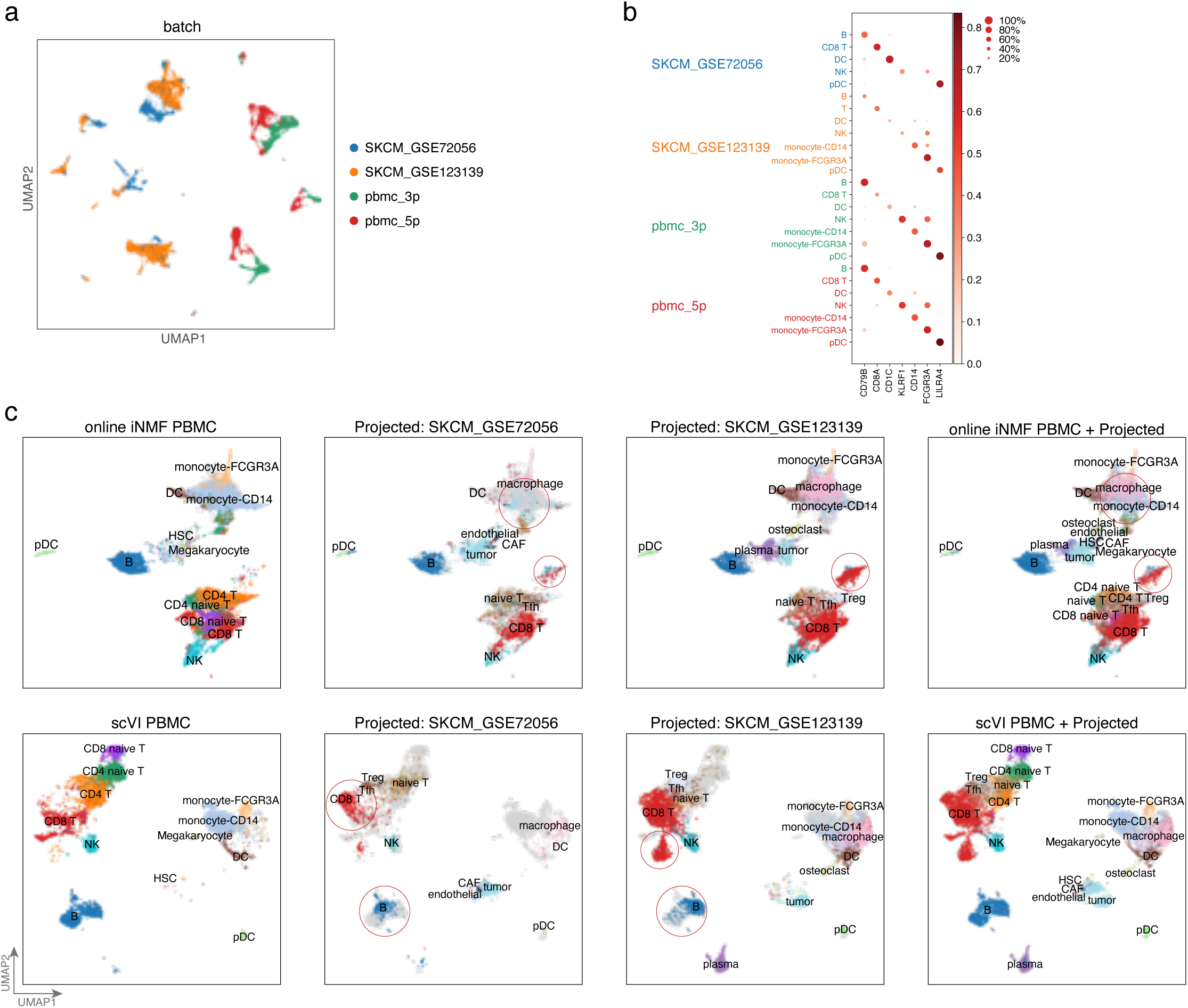
Projection of two melanoma datasets onto the *PBMC* dataset. **a**, UMAP embeddings of the *PBMC* dataset and the two additional melanoma data batches before integration. Cells are colored by batch. **b**, Dot plot of canonical markers of cell-types of each batch; dot color represents average expression level, while dot size represents the proportion of cells in the group expressing the marker. **c**, UMAP embeddings of the common cell space obtained by using online iNMF (top) and scVI (bottom) to project the two additional indicated melanoma data batches onto the *PBMC* dataset. Cells are colored by cell-type with light gray shadows representing the original *PBMC* dataset. Misalignments are highlighted with red circles. Note that here for convenient comparison of the projected data with the existing data, we showed the UMAP embeddings of the *PBMC* dataset combined with the projected data, which are visualized differently from the UMAP embeddings of the *PBMC* dataset alone in Fig. S2 as UMAP visualizations change with embedding data.

**Fig. S13.**
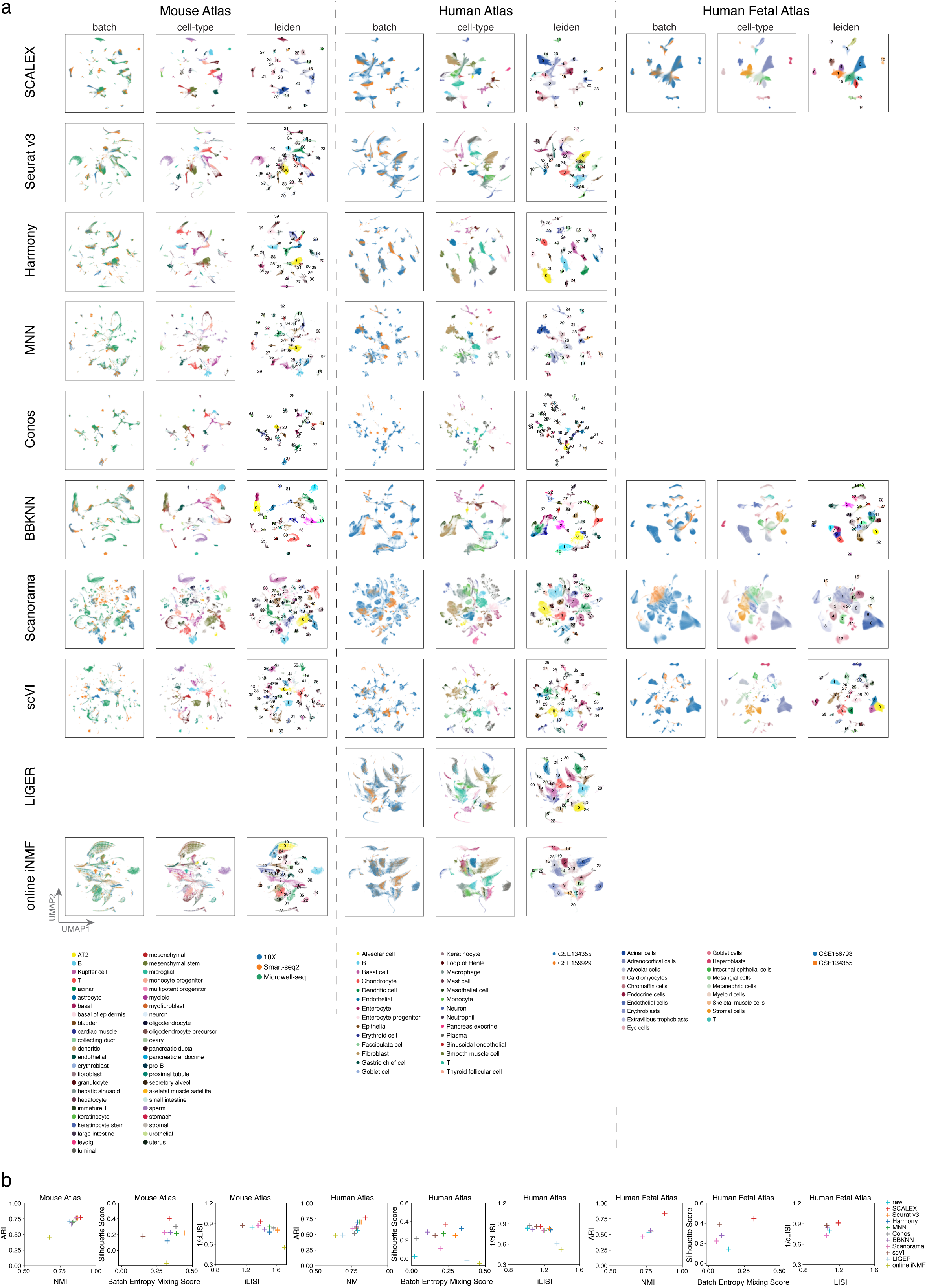
Comparisons of integration across Atlas-level datasets. **a**, UMAP embeddings of the indicated methods across the indicated datasets. Results are blank for those methods not scalable to this data size. **b**, Scatter plots showing the comparisons of indicated methods in terms of the ARI and NMI scores based-on the Louvain clustering results, the Silhouette and batch entropy mixing scores, and the cLISI/ iLISI scores based on the indicated Atlas-level datasets.

**Fig. S14.**
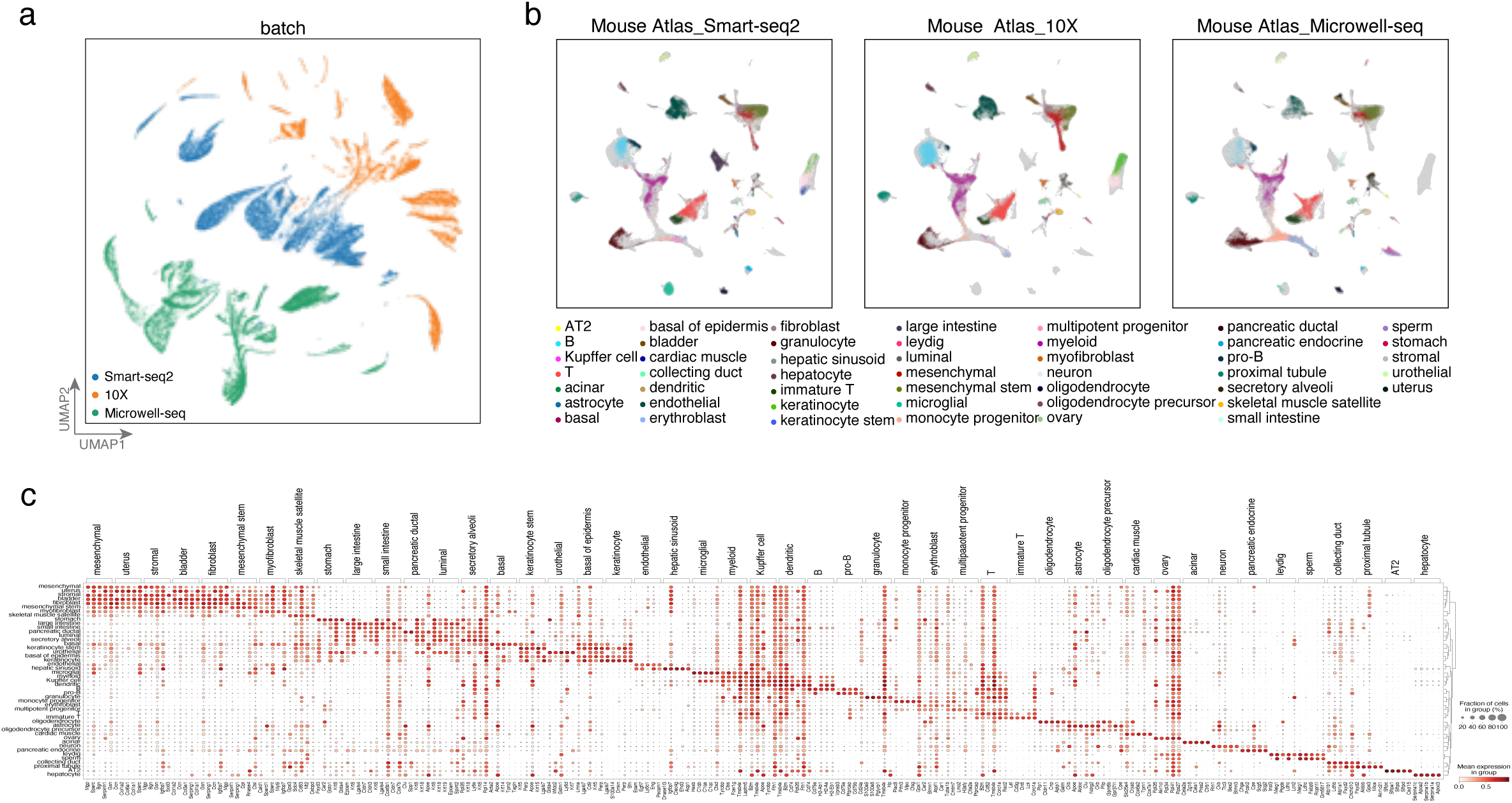
The SCALEX Mouse Atlas. **a**, UMAP embeddings of the Mouse Atlas dataset before integration, colored by batch. **b**, UMAP embeddings of three mouse atlas data batches (Smart-seq2, 10X, and Microwell-seq) after integration, colored by cell-type; the light gray shadows represent the original Mouse Atlas dataset. **c**, Dotplot of the top 5 cell-type-specific genes for each cell-type in the Mouse Atlas dataset. Dot color represents average expression level, while dot size represents the proportion of cells in the group expressing the marker. Note that here for convenient comparison of the projected data with the existing data, we showed the UMAP embeddings of the Mouse Atlas combined with the projected data, which are visualized differently from the UMAP embeddings of the Mouse Atlas alone in Fig. S13 as UMAP visualizations change with embedding data.

**Fig. S15.**
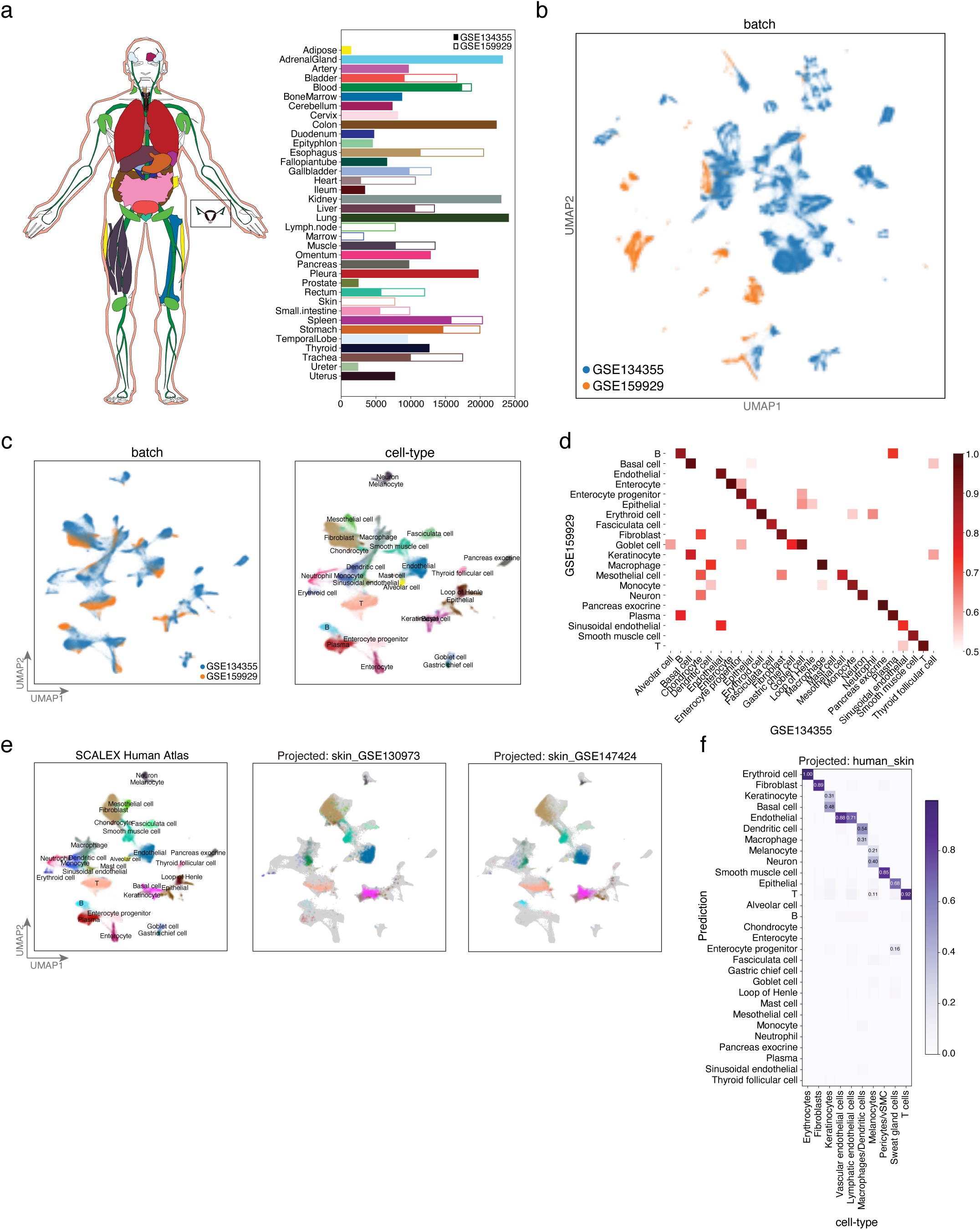
The SCALEX Human Atlas. **a,** The Human Atlas dataset acquired using different technologies (Smart-seq2, 10X, and Microwell-seq) covering various tissues used for construction of the human atlas. **b-c**, UMAP embeddings of the Human Atlas dataset colored by batch and cell-type, before (**b**) and after integration (**c**). **d**, Similarity matrix of meta-cell representations for cell-types in the two data batches in the common cell-embedding space after SCALEX integration between two batches. Color bar represents the Pearson correlation coefficient between the average meta-cell representation of two cell-types from a respective data batch. **e**, UMAP embeddings of the common cell space obtained by using SCALEX to project the two additional indicated human skin data batches onto the Human Atlas dataset. Cells are colored by cell-type with light gray shadows representing the original Human Atlas dataset. **f**, Confusion matrix of the cell-type annotations by SCALEX and those in the original study. Color bar represents the percentage of cells in confusion matrix *C_ij_* known to be in cell-type *i* and predicted to be in cell-type *j*. Note that here for convenient comparison of the projected data with the existing data, we showed the UMAP embeddings of the Human Atlas combined with the projected data, which are visualized differently from the UMAP embeddings of Human Atlas alone in Fig. S13 as UMAP visualizations change with embedding data.

**Fig. S16.**
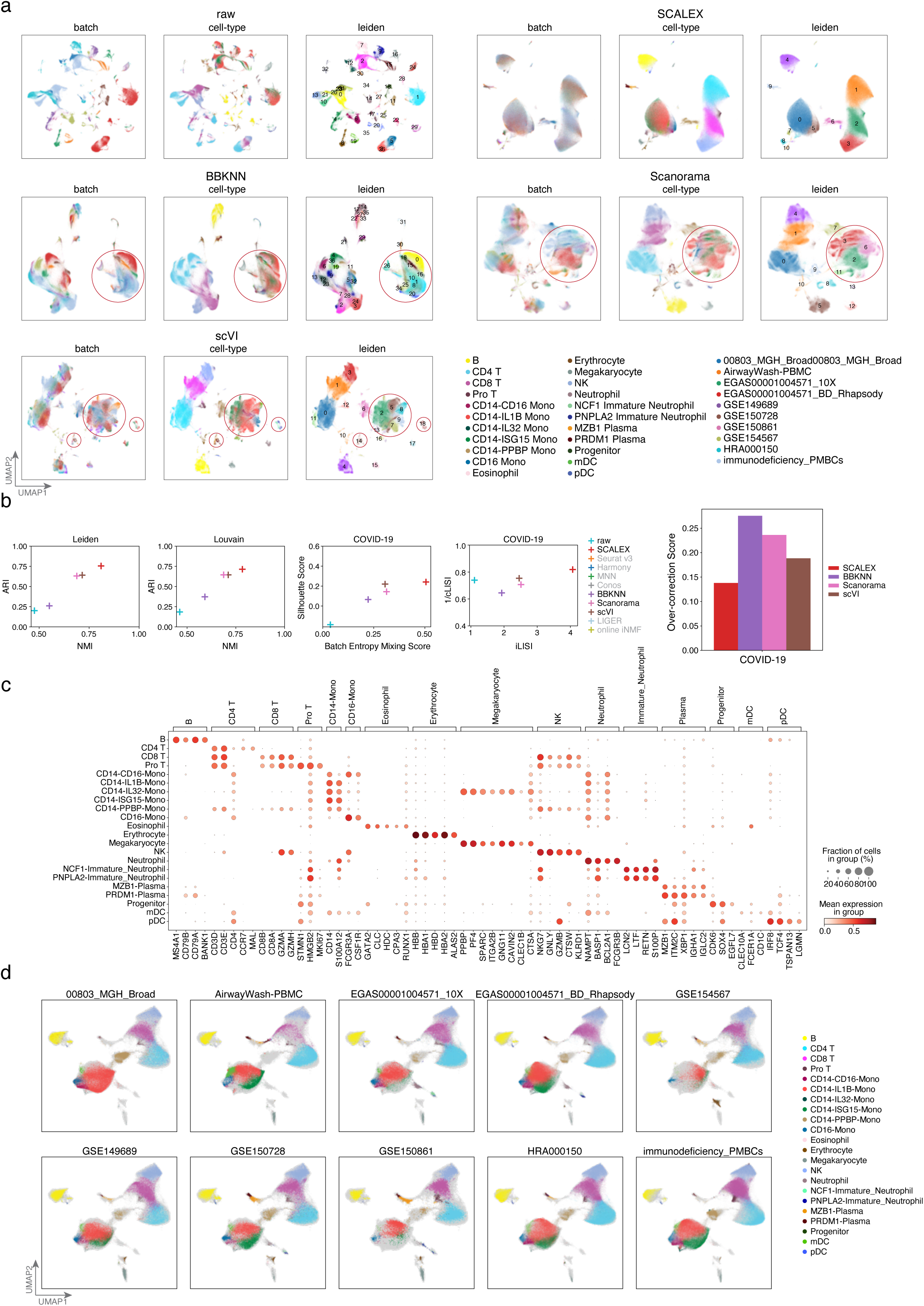
COVID-19 immune landscape. **a**, UMAP embeddings of the COVID-19 PBMC Atlas dataset before and after integration by indicated methods. Cells are colored by batch (left), cell-type (middle) and Leiden clustering results (right). **b**, Scatter plots showing the comparisons of indicated methods in terms of the ARI and NMI scores based-on the Leiden and Louvain clustering results, the Silhouette and batch entropy mixing scores, and the cLISI/ iLISI scores. Bar plot showing the comparisons of indicated methods in terms of the over-correction scores based on the COVID-19 PBMC Atlas dataset. **c**, Dotplot of canonical marker genes for each cell-type. Dot color represents average expression level, while dot size represents the proportion of cells in the group expressing the marker. **d**, UMAP embeddings of the COVID-19 PBMC atlas in individual batches after SCALEX integration, colored by cell-type; the light gray shadows represent the other batches of COVID-19 PBMC atlas. Note that here for convenient comparison of the projected data with the existing data, we showed the UMAP embeddings of the SCALEX COVID-19 PBMC Atlas combined with the projected data, which are visualized differently from the UMAP embeddings of SCALEX COVID-19 PBMC Atlas alone as UMAP visualizations change with embedding data.

**Fig. S17.**
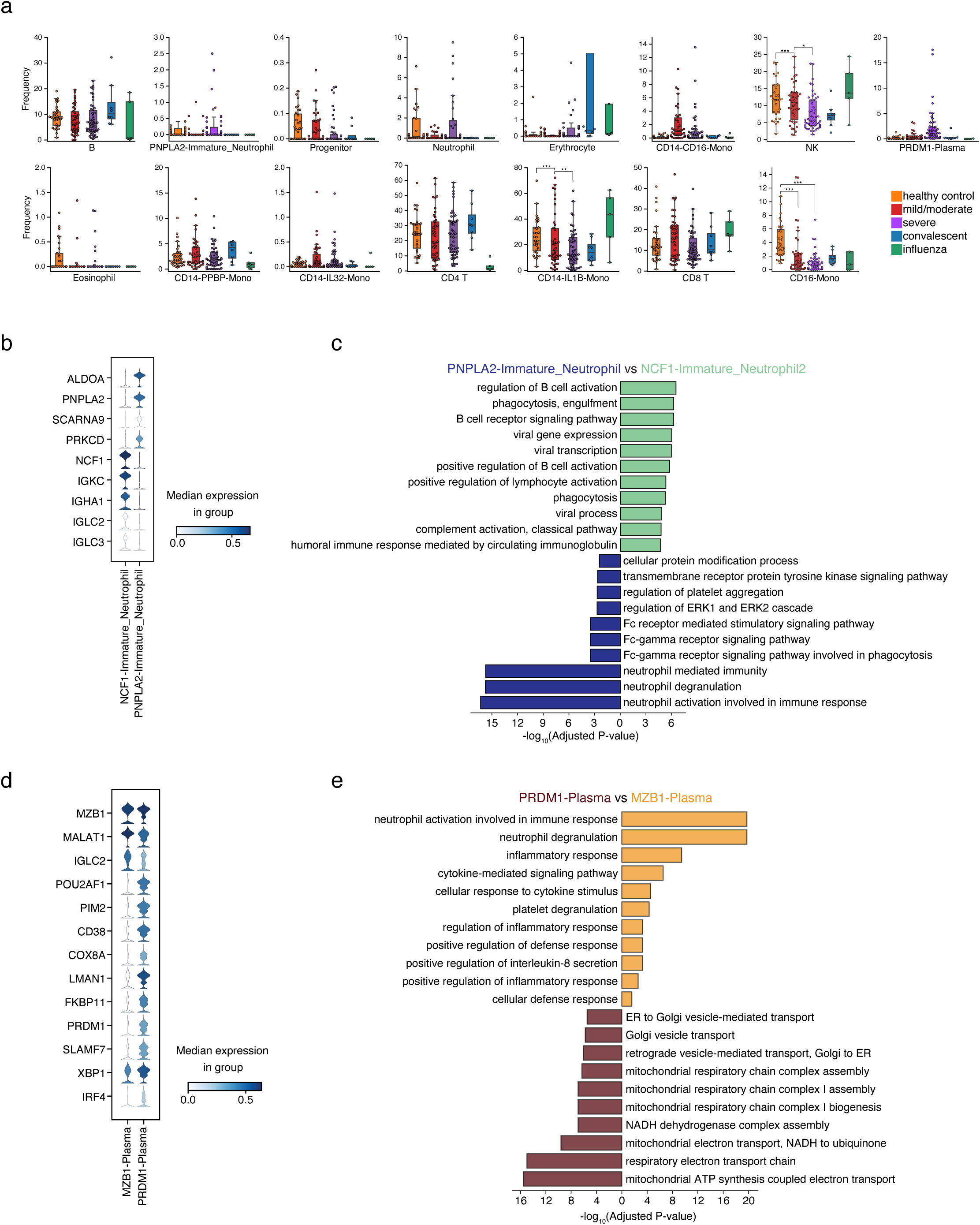
COVID-19 heterogeneous dysfunctional immune response. **a**, Frequency of cell distributions across healthy people and influenza patient controls, and among mild/moderate, severe, and convalescent COVID-19 patients. Dirichlet-multinomial regression was used for pairwise comparisons, ***p<0.001, **p<0.01, *p<0.05. **b**, Stacked violin plot of differentially-expressed genes between PNPLA2-Immature_Neutrophil and NCF1-Immature_Neutrophil cells. **c**, GO terms enriched in the differentially-expressed genes for PNPLA2-Immature_Neutrophil and NCF1-Immature_Neutrophil cells. **d**, Stacked violin plot of differentially-expressed genes between PRDM1-Plasma and MZB1-Plasma. **e**, GO terms enriched in the differentially-expressed genes for PRDM1-Plasma and MZB1-Plasma cells.

**Fig. S18.**
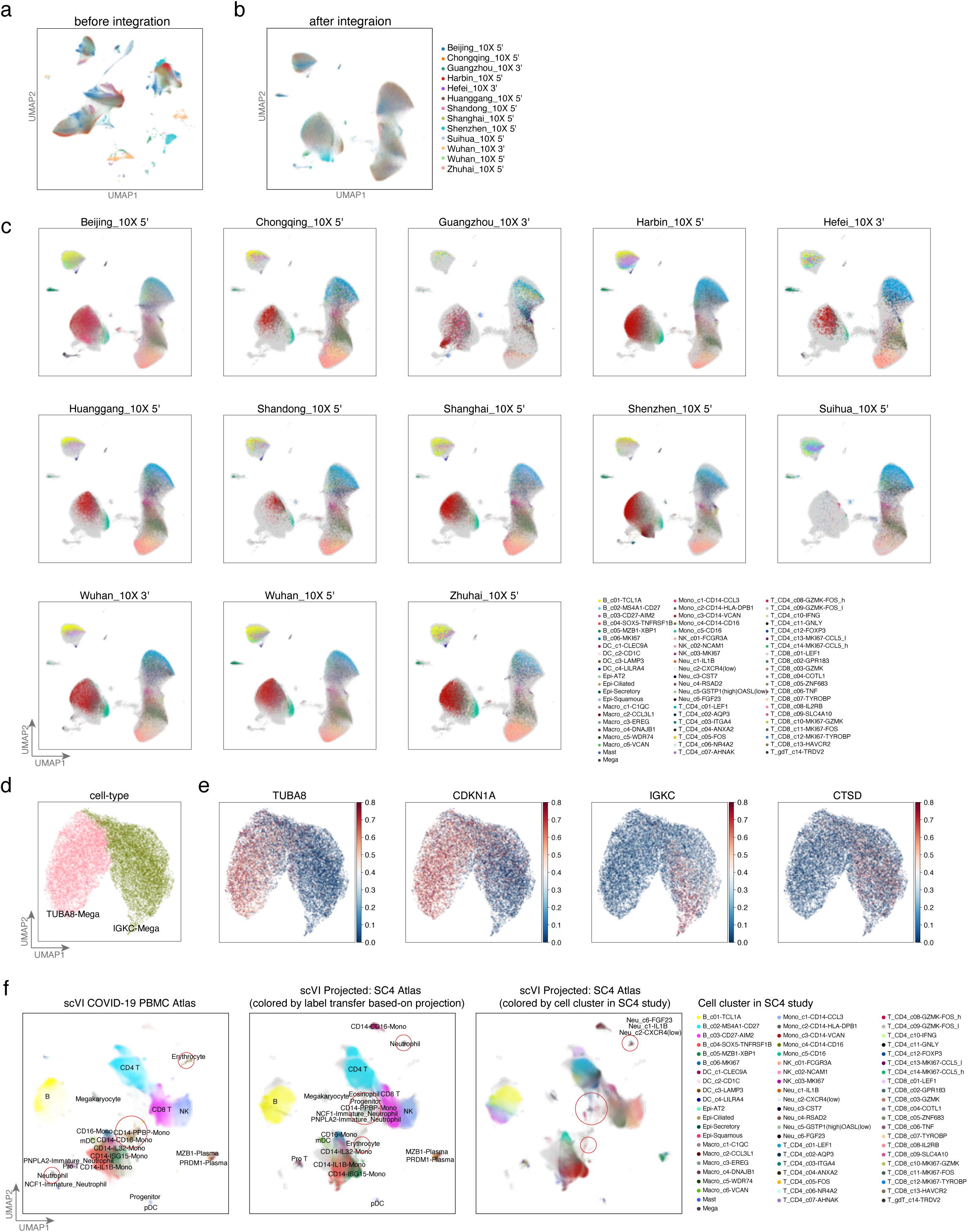
Projection of the SC4 Atlas onto the SCLAEX COVID-19 PBMC Atlas. **a-b**, UMAP embeddings of the SC4 Atlas before integration (**a**) and after projection onto the SCALEX COVID-19 PBMC Atlas (**b**). **c**, Separate UMAP embeddings of each SC4 data batch, after being projected onto the SCALEX COVID-19 PBMC space, colored by cell-type. light gray shadows represent the COVID-19 PBMC Atlas. **d**, UMAP embeddings of the TUBA8-Mega and IGKC-Mega cells. **e**, UMAP embeddings of the differentially-expressed genes of TUBA8-Mega and IGKC-Mega cells. **f**, UMAP embeddings of the common cell space obtained by using scVI to project the *SC4 Atlas* onto the COVID-19 PBMC Atlas dataset. Cells are colored by cell-type with light gray shadows representing the original COVID-19 PBMC Atlas dataset. Misalignments are highlighted with red circles.

**Fig. S19.**
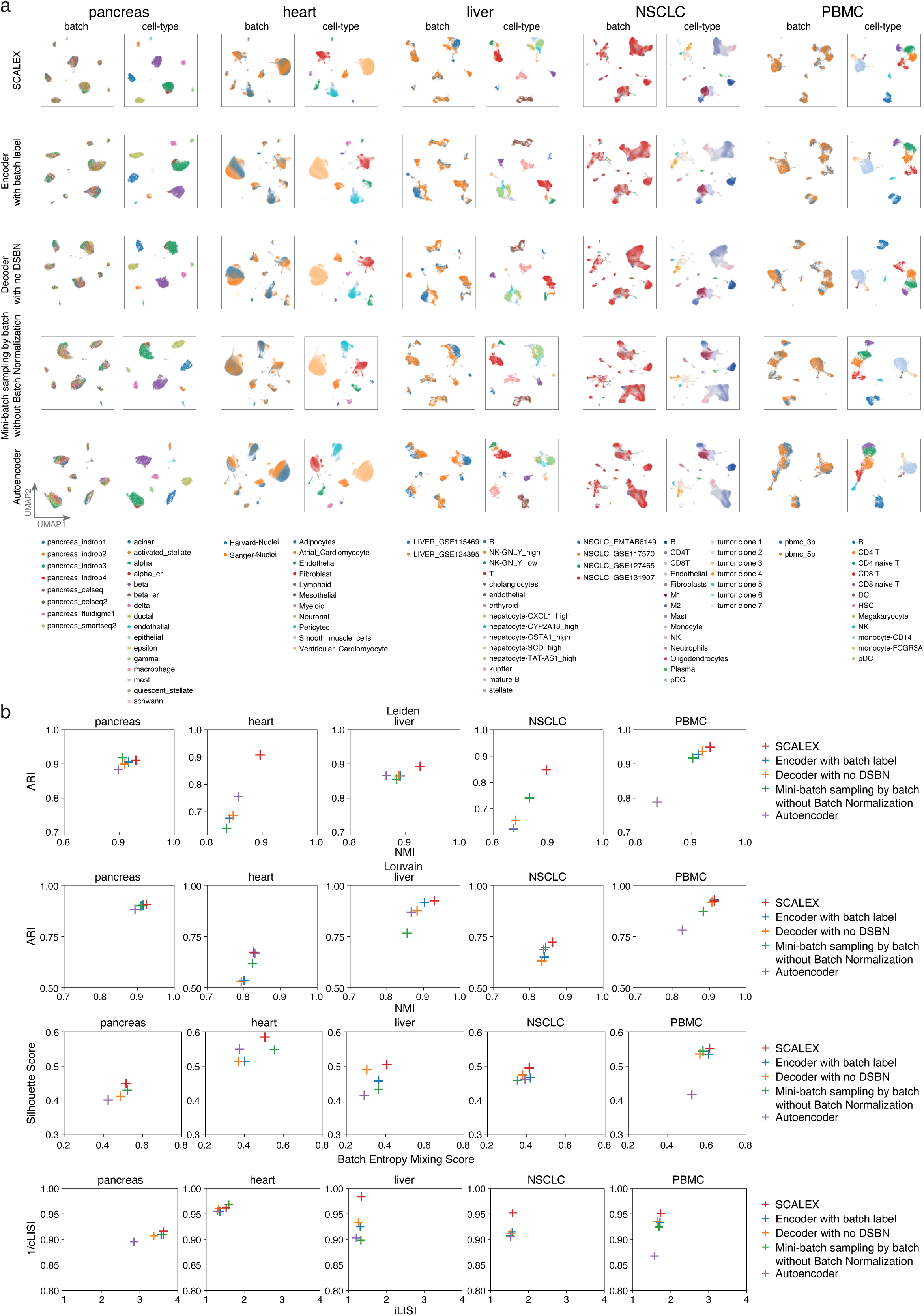
Ablation studies of different SCALE architectures for single-cell data integration. **a**, UMAP embeddings after integration by SCALEX and other SCALEX test-variants. **b**, Scatter plot showing a quantitative comparison of the performance of the full and different test-variants of SCALEX using the ARI score (y-axis) and the NMI score (x-axis), the Silhouette score (y-axis) and the batch entropy mixing score (x-axis), and the 1/cLISI score (y-axis) and the iLISI score (x-axis), across the indicated benchmark datasets.

**Fig. S20.**
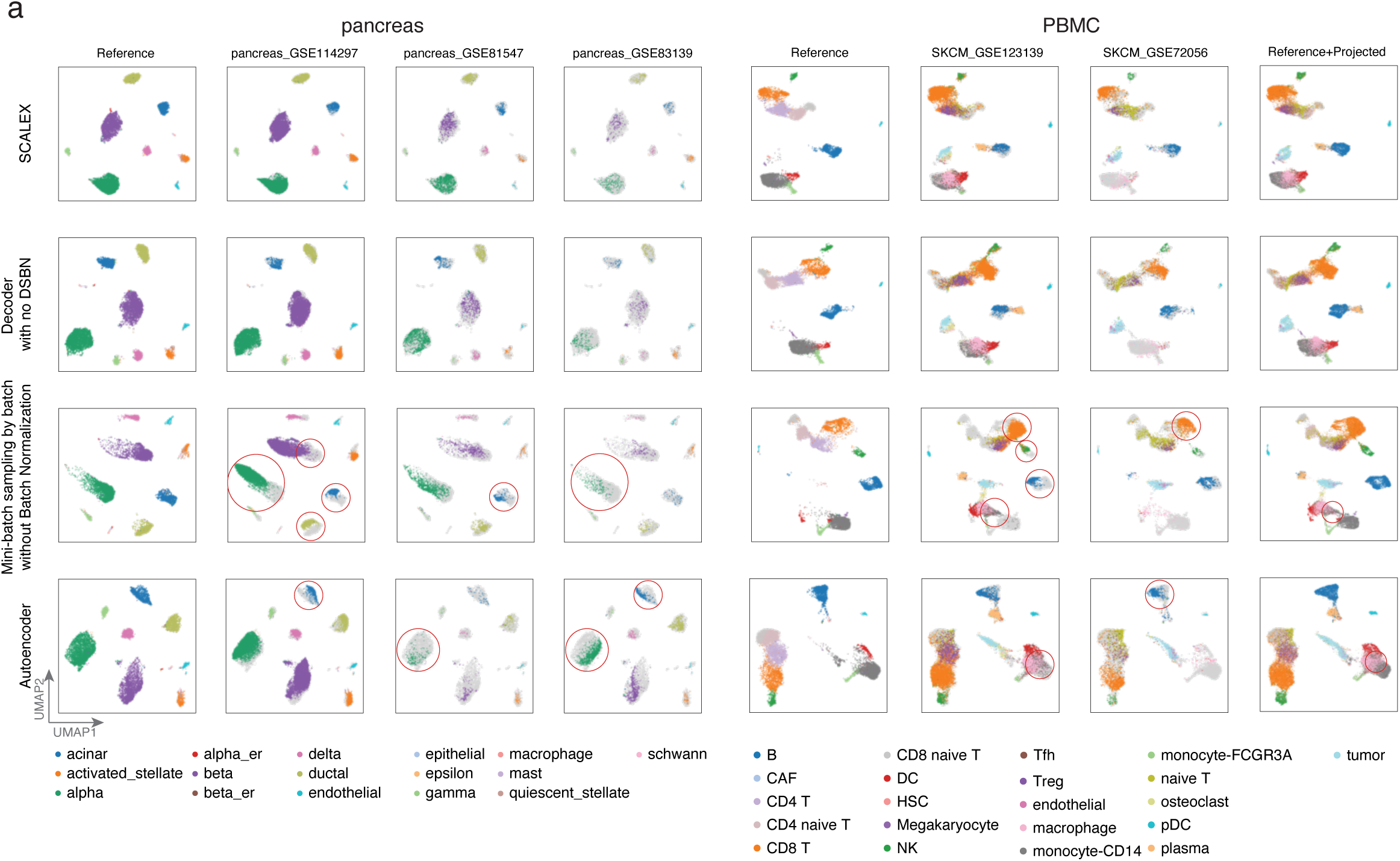
Ablations studies of different SCALEX architectures for single-cell data projection. Left: UMAP embeddings of three projected pancreas data batches projected onto the pancreas space using different SCALEX architectures, colored by cell-type; light gray shadows represent the original *pancreas* dataset. Right: UMAP embeddings of the two projected melanoma data batches projected onto the PBMC space using different SCALEX architectures, colored by cell-type; light gray shadows represent the original *PBMC* dataset.

**Fig. S21.**
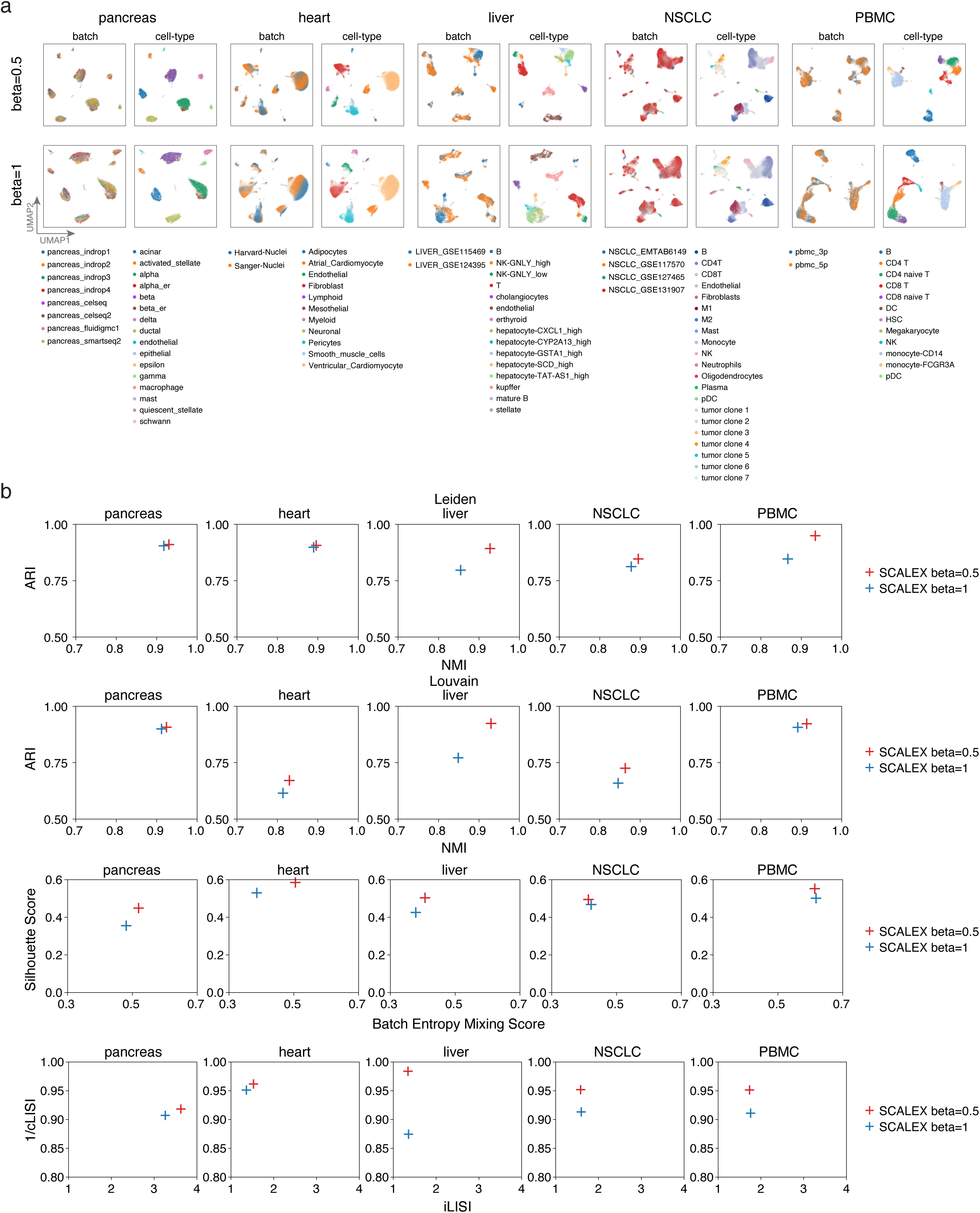
Comparison of SCALEX with beta=0.5 and beta=1 across the indicated benchmark datasets. **a**, UMAP embeddings of SCALEX (beta=0.5) and SCALEX (beta=1) across the indicated benchmark datasets; colored by batch (left) and cell-type (right). **b**, Scatter plot showing a quantitative comparison of the performance of SCALEX (beta=0.5) and SCALEX (beta=1) using the ARI score (y-axis) and the NMI score (x-axis), the Silhouette score (y-axis) and the batch entropy mixing score (x-axis), and the 1/cLISI score (y-axis) and the iLISI score (x-axis), across the indicated benchmark datasets.

## Supplementary Note: Ablation Study

To accomplish an accurate generalized encoder, the design of the full SCALEX included the following specific innovations:

1. an asymmetric autoencoder that inputs batch information only to the decoder (*i.e.*, never to the encoder) (See diagram in Fig. S1);
2. a DSBN layer in the decoder to release the encoder from the burden of capturing the batch-specific variations;
3. a mini-batching strategy that samples data from all batches simultaneously (rather than single batches iteratively) and thus more tightly follows the same overall distribution of the full input dataset; this strategy includes a Batch Normalization layer in the encoder that adjusts the deviation of each mini-batch and aligns them to the overall input distribution.

We conducted ablation studies to investigate the contributions of each design element of SCALEX. That is, we analyzed the performance (for integration and projection tasks) of the full SCALEX and four SCALEX “test-variants”, each with a distinct network architecture (Fig. S19-21). These can be summarized as follows:

### Full SCALEX (referred as “Baseline” in the following)

This Baseline model includes an encoder without batch labeling, sampling from all batches with Batch Normalization, decoder with DSBN, and beta=0.5. All other ablations are compared relative to this.

### Encoder with batch label

A test-variant with Baseline including an encoder with batch label as input. We found that the integration performance of this variant is similar with the full SCALEX, showing only a slight reduction in the evaluation scores (Fig. S19). However, the real issue is this: this addition of batch information at the beginning precludes online integration of newly arriving data. Put another way, this SCALEX test-variant is not capable of integrating single-cell data in a truly online manner.

### Decoder without DSBN

A test-variant of Baseline removing the DSBN layer from decoder. The DSBN layer combines multiple batch-specific Batch Normalization layers to capture the batch-specific information; this approach has been demonstrated as effective for domain adaption, as it provides a weak alignment across different domains. We observed an obvious drop in the integration performance (in terms of all evaluation scores) and a slight drop in the projection performance of this SCALEX test-variant, based-on the UMAP embeddings (Fig. S19,20).

### Sampling by batch without the Batch Normalization (BN) layer in the encoder

Removing the Batch Normalization layer from the encoder, and each mini-batch is sampled by batch instead of from all the batches. The integration performance of this test-variant was obviously worse than full SCALEX (Fig. S19). The projection performance also dropped obviously, with clear deviations from the common cell-embedding space (Fig. S20).

### Regular autoencoder

A test-variant that uses a regular autoencoder instead of a VAE framework. This variant performed the worst among all the variants we tested, for both the integration and projection tasks (Fig. S19,20).

Note that we also explored altering the Beta factor. Replacing the beta factor as 1 instead of 0.5. The integration performance of test variant of SCALEX for beta factor of 1 is worse than the SCALEX for beta factor of 1 (Fig. S21).

